# Versatile multi-transgene expression using improved BAC TG-EMBED toolkit, novel BAC episomes, and BAC-MAGIC

**DOI:** 10.1101/708024

**Authors:** Binhui Zhao, Pankaj Chaturvedi, David L. Zimmerman, Andrew S. Belmont

**Affiliations:** Department of Cell and Developmental Biology, University of Illinois, Urbana, Illinois, 61801, United States of America

**Author notes:** The authors wish it to be known that, in their opinion, the first two authors should be regarded as joint First Authors. David L. Zimmerman, Biology Department, College of the Ozarks, Point Lookout, Missouri, 65726, United States of America.

## Abstract

Achieving reproducible, stable, and high-level transgene expression in mammalian cells remains problematic. Previously, we attained copy-number-dependent, chromosome-position-independent expression of reporter minigenes by embedding them within a BAC containing the mouse *Msh3*-*Dhfr* locus (DHFR BAC). Here we extend this “BAC TG-EMBED” approach. First, we report a toolkit of endogenous promoters capable of driving transgene expression over a 0.01-5 fold expression range relative to the CMV promoter, allowing fine-tuning of relative expression levels of multiple reporter genes expressed on a single BAC. Second, we show small variability in both the expression level and long-term expression stability of a reporter gene embedded in BACs containing either transcriptionally active or inactive genomic regions, making choice of BACs more flexible. Third, we describe an intriguing phenomenon in which BAC transgenes are maintained as episomes in a large fraction of stably selected clones. Finally, we demonstrate the utility of BAC TG-EMBED by simultaneously labeling three nuclear compartments in 94% of stable clones using a multi-reporter DHFR BAC, constructed with a combination of synthetic biology and BAC recombineering tools. Our extended BAC TG-EMBED method provides a versatile platform for achieving reproducible, stable simultaneous expression of multiple transgenes maintained either as episomes or stably integrated copies.

## INTRODUCTION

Transgene expression has been widely used in both basic research and biotechnology. Applications of transgene expression range from the elucidation of gene function by ectopic expression of selected transgenes, to the expression of transgenes for gene therapy, and to the overexpression of genes for production of biopharmaceuticals (1–5). Examples of such applications include the expression of multiple fluorescent proteins for live-cell imaging (6), the expression of the four or more Yamanaka transcription factors for efficient generation of induced pluripotent stem (iPS) cells (7), and the expression of multiple proteins for reconstitution of protein complexes (8).

Despite the currently widespread use of transgene expression, most transgene expression systems still suffer from serious experimental limitations. Plasmid-, lentivirus- and transposon-based systems, all still show varying degrees of chromosome position effects (9, 10) and position effect variegation (PEV) (11–15). Moreover, foreign sequences by themselves are targets for epigenetic silencing (16–19), and transgene concatamers can induce the formation of heterochromatin (20, 21). Together these transgene silencing mechanisms result in unpredictable transgene expression levels that do not correlate with copy number and are unstable with long-term culture or changes in the cell physiological or differentiated state (22–24).

Such limitations are compounded when the simultaneous and reproducible expression of multiple transgenes is required. For example, a common application in the emerging field of synthetic biology is the design of novel gene circuits, involving the expression of multiple proteins, in many cases at precise relative levels (25). While this approach has worked well in prokaryotes and yeast, it has been difficult to implement in mammalian cells due to the lack of suitable multi-transgene expression methods which overcome chromosome position effects and allow expression of different transgenes at reproducible relative levels.

A commonly used approach to countering transgene silencing and variegation has been through the inclusion of *cis*-elements. These include insulators (26, 27), locus control regions (LCRs) (28, 29), scaffold/matrix attachment regions (S/MARs) (30, 31), ubiquitous chromatin opening elements (UCOEs) (32, 33) and anti-repressors (34); some of these regulatory elements have context-dependent and/or vector dependent activity. While these *cis*-elements improve transgene expression to varying degrees, they are insufficient for chromosome-position independent, copy-number-dependent transgene expression (29, 35–37).

Additionally, in some transgene expression applications the ability to avoid transgene chromosomal integration and eventually eliminate these transgenes from the cells is highly desirable. Both viral-sequence based and non-viral, pEPI based episomal vectors have been developed (38–41). Viral-based vectors have the potential of causing transformation of the transfected cells (42), while pEPI-like vectors, containing a S/MAR sequence immediately downstream of an active transcription unit, are mitotically stable without selection (43–47), and thus cannot be removed from the cells. Moreover, transgenes on these episomal vectors are still subject to silencing (48), possibly due to the prokaryotic or viral sequences on these vectors (49, 50).

Bacterial artificial chromosomes (BACs) carrying ∼100-200 kb mammalian genomic DNA inserts harbor most of the *cis*-regulatory sequences required for expression of the endogenous genes contained within these genomic inserts. Previously we demonstrated how embedding minigene constructs at different locations within the DHFR BAC provided reproducible expression of single or multiple reporter genes independent of the chromosome integration site (51). Similar approaches were used by other labs for high-level recombinant protein production (52, 53). Recently, our lab demonstrated stable transgene expression after cell-cycle arrest or after terminal cell differentiation, using the BAC-TG EMBED approach (54). All of these studies tested only BACs containing actively transcribed regions, based on the hypothesis that the expression level of the transgenes inserted into the BACs was determined by the chromatin environments reconstituted by the genomic inserts within the BACs. Indeed, because of this assumption, previous studies have specifically targeted the inserted transgenes to transcription units and even exons (51–53).

However, this hypothesis has not been tested. Moreover, overexpression from the genes on the BAC genomic inserts might change the properties of the transfected cells, or interfere with other assays of a study. Thus, BACs with no transcription units would be more desirable. Another improvement over our previous BAC TG-EMBED system (51, 54) would be a toolkit of endogenous promoters capable of driving transgene expression over a wide range of defined, relative expression levels. Viral promoters, including the CMV promoter we used previously, are known to be prone to epigenetic silencing (55, 56), while most previously used endogenous and synthetic promoters were selected for their strength (53, 57–60). While high-level transgene expression is preferable in applications calling for overexpression, a low or near-physiological expression is important for many other applications, including gene therapy. Additionally, multiple transgenes may need to be expressed simultaneously but at reproducible differential levels.

Here we describe further extensions to the BAC TG-EMBED method that together provide a more versatile BC TG-EMBED toolkit for a range of future potential applications. First, we describe a toolkit of endogenous promoters, for which we have measured relative promoter strength, that can drive transgene expression at reproducible relative levels over a 500-fold range. Second, we show that multiple BAC scaffolds can be used to drive sustained high-level transgene expression driven by the UBC promoter without selection for up to 12 weeks, including BAC scaffolds containing no active transcription units. Third, we describe an episomal version of BAC TG-EMBED, where BAC transgenes form circular, ∼1 Mb episomes and can be eliminated from the cells by removing selection. Fourth, we developed a “BAC-MAGIC” (**<u>BAC</u>**-**<u>M</u>**odular **<u>A</u>**ssembly of **<u>G</u>**enomic loci **I**nterspersed **<u>C</u>**assettes) to more rapidly assemble BACs containing multiple transgene expression cassettes. Finally, as a proof-of-principle demonstration of our new, more versatile BAC TG-EMBED toolkit, we demonstrate simultaneous expression of fluorescently tagged proteins labeling three different nuclear compartments, achieving >90% optimally labeled cell clones after a single, stable transfection.

## MATERIALS AND METHODS

### PCR amplification of endogenous promoters

Primers (Supplementary Table S1) were designed using Primer3 (61) or NCBI primer blast (62) to amplify 1-3 kb promoter regions which included either the entire or part of the 5’ UTRs upstream of the first exons of target genes. We used human genomic DNA extracted from BJ-hTERT cells as the template for PCR. However, the UBC promoter, including a partially synthetic intron, was amplified from plasmid pUGG (54).

### Construction of dual reporter DHFR BACs

The original dual reporter BAC, DHFR-HB1-GN-HB2-RZ (51), was derived from the CITB-057L22 BAC (DHFR BAC) containing mouse chr13:92992156-93161185 (mm9). DHFR-HB1-GN-HB2-RZ has an EGFP expression cassette inserted 26 kb downstream of the *Msh3* transcription start site, and a mRFP expression cassette inserted at 121 kb downstream of the *Msh3* transcription start site. The EGFP expression cassette contains a CMV promoter-driven EGFP gene and a SV40 promoter-driven Kanamycin/Neomycin resistance gene, while the mRFP expression cassette has a CMV promoter-driven mRFP gene and a SV40-driven Zeocin resistance gene. New dual reporter DHFR BACs were created using a similar strategy to that used to create DHFR-HB1-GN-HB2-RZ, except that new mRFP expression cassettes were used, where the CMV promoter was replaced with alternative, human endogenous promoters. The intermediate DHFR BAC containing only the EGFP expression cassette, DHFR-HB1-GN (51), was used to insert these new mRFP expression cassettes using *λ* Red-mediated homologous recombination (63, 64).

Plasmid p[MOD-HB2-CRZ] (51) contains a CMV driven mRFP and a SV40 driven Zeocin resistance gene, flanked by two ∼500 bp regions homologous to the DHFR BAC target site. Plasmid p[MOD-HB2-RCS-Zeo] was created by replacing the CMV-mRFP fragment between NotI and NheI sites of p[MOD-HB2-CRZ] with a synthetic DNA fragment “RCS” containing multiple rare restriction sites (Supplementary Table S2). The mRFP fragment generated by digesting p[MOD-HB2-CRZ] with NheI was then inserted into the NheI site of p[MOD-HB2-RCS-Zeo], yielding plasmid p[MOD-HB2-RCS-RZ]. The PCR-amplified endogenous promoters were then inserted into the RCS, generating plasmids p[MOD-HB2-promoter name-RZ]. Promoter functionality was tested by transient transfection of NIH 3T3 cells with these plasmids.

To insert the new mRFP expression cassettes into the DHFR-HB1-GN BAC, one round of *λ* Red-mediated recombination, using Zeocin resistance as positive selection, was performed according to a published protocol (63). DNA fragments containing the new mRFP expression cassettes with a given promoter with flanking homologous arms were excised from p[MOD-HB2-promoter name-RZ] plasmids by PmeI. SW102, a derivative strain of *Escherichia coli* (*E. coli*), was used for recombination. Recombinants were selected on low-salt LB plates containing 25 µg/ml Zeocin and 12.5 µg/ml Kanamycin at 32°C for ∼20 hours. Recombinant colonies were screened by PCR amplification of sequences flanking the site of insertion (primers listed in Supplementary Table S1). The integrity of BAC constructs was verified by restriction enzyme fingerprinting, where observed band patterns on agarose gels were compared with predicted ones.

### Construction of BACs containing the UBC-GFP-ZeoR cassette

Construction of pUGG containing the UBC-GFP-ZeoR-FRT-GalK-FRT cassette was described previously (54). Human BACs RP11-138I1 (UBB BAC), CTD-2643I7 (HBB BAC), CTD-2207K13 (2207K13 BAC) and mouse BAC RP23-401D9 (ROSA BAC) were obtained from Thermo Fisher Scientific. Mouse BAC CITB-057L22 (DHFR BAC) was a gift from Edith Heard (Curie Institute, Paris, France).

The UBC-GFP-ZeoR reporter gene insertion positions (mm9 or hg19) are chr17:16,301,887-16,301,888 in the UBB BAC, chr6:113,043,332-113,043,333 in the ROSA BAC, chr13:93,099,101-93,099,102 in the DHFR BAC, chr1:79,224,725-79,224,726 in the 2207K13 BAC, and chr11:5,390,233-5,390,244 in the HBB BAC.

*λ* Red-mediated BAC recombineering (63, 64) using a *galK*-based dual-selection scheme was used to introduce the UBC-GFP-ZeoR reporter cassette onto the BACs according to published protocols (63). DNA fragments with homology ends for recombineering were prepared by PCR using primers (Supplementary Table S1) with 74-bp homology sequences plus 16-bp sequences (forward, 5’-acagcagagatccagt-3’; reverse, 5’-tgttggctagtgcgt-3’) that amplify the UBC-GFP-ZeoR-FRT-GalK-FRT cassette from plasmid pUGG. *E. coli* strain SW105 was used for BAC recombineering. Recombinants containing the UBC-GFP-ZeoR-FRT-GalK-FRT cassette were selected for *galK* insertion at 32°C on minimal medium in which D-galactose was supplied as the only carbon source. Recombinant colonies were screened using PCR with BAC specific primers flanking the target regions (Supplementary Table S1). Subsequently, FLP recombinase-mediated removal of *galK* from selected recombinant clones was done by inducing actively growing SW105 cells with 0.1% (w/v) L-arabinose. Negative selection against *galK* used minimal medium containing 2-deoxy-galactose; deletion of *galK* in recombinants was again verified using BAC specific primers (Supplementary Table S1). The integrity of BAC constructs was verified by restriction enzyme fingerprinting.

The UBB, HBB, 2207K13, ROSA, DHFR BACs with the UBC-GFP-ZeoR reporter gene inserted were named UBB-UG, HBB-UG, 2207K13-UG, ROSA-UG and DHFR-UG, respectively.

### Cell culture and establishment of BAC cell lines

Mouse NIH 3T3 fibroblasts (ATCC CRL-1658^TM^) were grown in Dulbecco’s modified Eagle medium (DMEM, with 4.5 g/l D-glucose, 4 mM L-glutamine, 1 mM sodium pyruvate and 3.7 g/l NaHCO_3_) supplemented with 10% HyClone Bovine Growth Serum (GE Healthcare Life Sciences, Cat. # SH30541.03). Human HCT116 cells (ATCC CCL-247^TM^) were grown in McCoy’s 5A medium supplemented with 10% Fetal Bovine Serum (Seradigm, Cat. # 1500-500H).

BAC DNA for transfection of mammalian cells was prepared with the QIAGEN Large Construct Kit (QIAGEN, Cat. # 12462) as per the manufacturer’s instructions. All BACs except DHFR BAC derived BACs were linearized before transfection: 2207K13-UG BAC with SgrAI (New England Biolabs, Cat. # R0603S), HBB-UG BAC with NotI (New England Biolabs, Cat. # R3189S) and all other BACs with the PI-SceI (New England Biolabs, Cat. # R0696S). Lipofectamine 2000 (Thermo Fisher Scientific, Cat. # 11668019) was used to transfect the cells with the BACs according to the manufacturer’s directions. The dual reporter DHFR BACs and the BACs containing the UBC-GFP-ZeoR reporter gene were transfected into NIH 3T3. The 2207K13-UG BAC was also transfected into HCT116. The DHFR BACs containing the Lac operator repeats were transfected into an NIH 3T3 cell clone 3T3_LG_C29 stably expressing the EGFP-dimer LacI-NLS fusion protein (EGFP-LacI) (65). Mixed clonal populations of stable transformants were obtained after ∼2 weeks of selection (75 µg/ml Zeocin and 500 µg/ml G418 for NIH 3T3 cells transfected with the dual reporter DHFR BACs; 75 µg/ml or 200 µg/ml Zeocin for NIH 3T3 or HCT116 cells, respectively, transfected with the BACs containing the UBC-GFP-ZeoR reporter gene; 75 µg/ml Zeocin and 200 µg/ml Hygromycin B for 3T3_LG_C29 transfected with the DHFR BACs); individual cell clones were obtained by serial dilution or colony picking using filter discs (66).

To analyze the stability of reporter gene expression in NIH 3T3 cells, individual cell clones were grown continuously with or without Zeocin (75 µg/ml) selection for 96 days. We used the following clones (Figure 4 and Supplementary Figure S1): DHFR-UG BAC-f1-7, f3-13, f3-15 (uniform), f1-6, f2-1, f2-3 (heterogeneous); ROSA-UG BAC-2D6-3C11, 3D7 (uniform), 2C12, 3A1 (heterogeneous); UBB-UG BAC-1C2, 1F1, 1F12, 2F5, 2G4, 4D3, 5C1, 5C7 (uniform), 1A8, 1D5, 6H2 (heterogeneous); 2207K13-UG BAC-3E3, 5C8, 5E1, 6B9, 6E12, 6F4, 7B2 (uniform), 1E3, 6A2, 6C10, 7B9 (heterogeneous).

### Flow cytometry

For analysis of reporter gene expression, cells were grown to ∼40%-80% confluence, trypsinized, and resuspended in growth media at ∼0.5-1 million/ml. For analysis of the expression of mRFP and EGFP, or mRFP alone, cell suspensions were run on a BD FACS AriaII (BD Biosciences) or a BD LSR Fortessa (BD Biosciences), using the PE channel (561 nm laser and 582/15 nm bandpass filter) for mRFP, and the FITC channel (488 nm laser, 505 longpass dichroic mirror and 530/30 nm bandpass filter) for EGFP. For analysis of GFP expression alone, the cell suspensions were run on a BD FACS Canto II Flow Cytometry Analyzer (BD Biosciences), using the FITC/Alexa Fluor-488 channel (488nm laser, 502 longpass dichroic mirror and 530/30 bandpass filter). Rainbow fluorescent beads (Spherotech, Cat. # RFP-30-5A) were used as fluorescence intensity standards. Each sample was run for 1-2 min or until the number of events after gating reached 10-20 thousand.

For cell sorting, cells were resuspended at ∼10 million/ml in growth media and run on a BD FACS AriaII for up to 30-40 minutes. Sorting windows are shown in the main and supplementary figures.

### Estimation of relative promoter strength

The red and green fluorescence of the mixed-clonal populations stably transfected with the dual-reporter DHFR BACs was measured by flow cytometry. The mean florescence values of all gated cells were divided by the bead intensity values for normalization. The ratio of normalized mRFP to normalized EGFP was calculated as a measure of promoter strength (Equation 1). All promoter strengths were then normalized with the CMV promoter strength (comparing the CMV-driven mRFP to the CMV-driven EGFP expression) to calculate the relative promoter strength (Equation 2) using the CMV promoter as the reference.

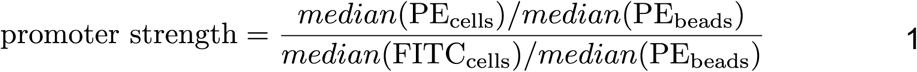

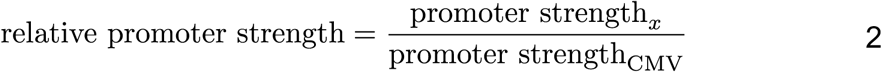

### Genomic DNA extraction

Genomic DNA was isolated by phenol/chloroform extraction (67). Cultured cells were harvested and washed with 1x Cell Culture Phosphate Buffered Saline (PBS, Corning, Cat. # 21040CV). Sorted cells were pelleted. Up to ∼2 million cells were resuspended in 100 μl High-TE buffer (10 mM Tris-Cl, pH 8, 10 mM EDTA, 25-100 μg/ml RNase A (QIAGEN, Cat. # 19101)) and lysed by adding 2.5 μl 20% SDS. After incubation at 37°C for several hours, the lysate was digested by ∼0.2 mg/ml Proteinase K (New England Biolabs, Cat. # P8102 or P8107S) at 55°C for ∼1 day. 1 M Tris-Cl (pH 8.0), 5 M NaCl and nuclease free water were added to the lysate to bring up the total volume to ∼600 μl and final concentrations of Tris-Cl to ∼0.1 M and NaCl to ∼0.2 M. The lysate was then extracted once with an equal volume of phenol/chloroform/isoamyl alcohol (25:24:1 mixture, Fisher Scientific, Cat. # BP1752I-400) and once with an equal volume of chloroform/isoamyl alcohol (24:1 mixture, MilliporeSigma, Cat. # C0549). DNA was precipitated by adding 2.5 volumes of 100% ethanol, washed with 70% ethanol and resuspended in EB (10mM Tris-Cl, pH 8.5).

### Estimation of transgene copy number

BAC or plasmid transgene copy number within individual cell clones or sorted cells was measured by real-time quantitative PCR (qPCR), using purified genomic DNA, iTaq universal SYBR Green Supermix (Bio-Rad Laboratories, Cat. # 1725121) and a StepOnePlus (Applied Biosystems). Relative quantitation methods were used for copy number calculation. Primers used for qPCR are listed in Supplementary Table S1. Mouse genes *Sgk1* and *Hprt1* were used as endogenous controls, assuming four copies of each gene per cell in NIH 3T3. For Figure 3d and Figure 5c, a primer pair (Zeo-GFP2for/rev) that binds to the UBC-GFP-ZeoR region was used to estimate transgene copy number. For Table 2, Table 3 and Supplementary Figure S5, in addition to Zeo-GFP2for/rev, 4 primer pairs binding to the DHFR BAC or 6 primer pairs binding to the HBB BAC were used to estimate the copy number of DHFR-UG or HBB-UG BAC, respectively. The ΔC_T_ method (Equations 3 and 5) was used to estimate the copy numbers of the PCR amplification regions on the UBC-GFP-ZeoR reporter gene or on the HBB BAC, and ΔΔC_T_ method (Equations 4 and 6) was used to estimate the copy numbers of the PCR amplification regions on the DHFR BAC. When multiple primer pairs were used for a region, the mean copy number of all PCR amplification regions was calculated as the copy number of that region. Equations 3 and 7 were used to calculate the fold increase of BAC copy numbers in H1 and H2 samples relative to L.

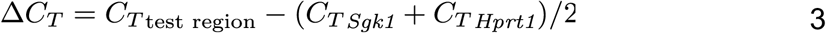

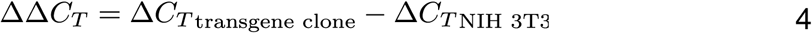

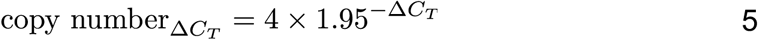

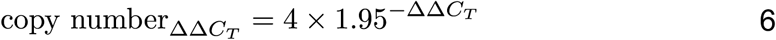

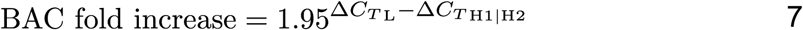

### Correlation of reporter gene expression and reporter gene copy number

Mean fluorescence intensity (in arbitrary units) of individual clones were measured by flow cytometry and normalized by fluorescent bead intensity to be used as a measure of reporter gene expression. To ensure uniform normalization for all samples, fluorescent beads from the same batch were used for all measurements. Untransfected cells were used to establish background fluorescence levels. Linear correlations of GFP expression level versus transgene copy number for each group of cell clones were calculated using the linear trend line tool in Microsoft Excel with the y-intercept fixed to 0 (autofluorescence normalized by beads was almost 0).

### DNA FISH probes

Biotin or digoxigenin labeled DNA FISH probes were made from BAC DNA, using a published protocol (68), with the following reagents: AluI, DpnI, HaeIII, MseI, MspI, RsaI (New England Biolabs, Cat. # R0137S, R0176S, R0108S, R0525S, R0106S, R0167S, respectively) and CutSmart Buffer (New England Biolabs); Terminal Deoxynucleotidyl Transferase and reaction buffer (Thermo Fisher Scientific, Cat. # EP0161); dATP (New England Biolabs, Cat. # N0446S) and Biotin-14-dATP (Thermo Fisher Scientific, Cat. # 19524016) for biotin labelling, or dTTP (New England Biolabs, Cat. # N0446S) and Digoxigenin-11-dUTP (MilliporeSigma, Cat. # 11093088910) for digoxigenin labelling.

### 3D DNA FISH

DNA FISH of interphase nuclei used published protocols (69, 70) with small modifications. Cells grown on coverslips (12 mm diameter) were fixed with 3-4% paraformaldehyde in Dulbecco’s phosphate buffered saline (DPBS, 8 g/l NaCl, 0.2 g/l KCl, 2.16 g/l Na_2_HPO_4_-7H_2_O, 0.2 g/l KH_2_PO_4_) for 10 min, followed by permeabilization with 0.5% Triton X-100 (Thermo Fisher Scientific, Cat. # 28314) in DPBS for 10-15 min. Cells were subjected to six freeze-thaw cycles using liquid nitrogen, immersed in 0.1M HCl for 10-15 min, and then washed 3x with 2x saline-sodium citrate (SSC). Freeze-thaw cycles sometimes were skipped with no noticeable difference in FISH signals. Cells were incubated in 50% deionized formamide (MilliporeSigma, Cat. # S4117)/2x SSC for 30 min at room temperature (RT), and stored for up to 1 month at 4°C. Each coverslip used ∼4 μl hybridization mixture, consisted of 5-20 ng/µl probes, 10x of mouse (for NIH 3T3 cells) or human (for HCT116 cells) Cot-1 DNA (Thermo Fisher Scientific Cat. # 18440016 or 15279011,) per ng probe, 50% deionized formamide, 10% dextran sulfate (MilliporeSigma, Cat. # D8906) and 2x SSC. Cells and probes were denatured together on a heat block at ∼76°C for 2-3 min and hybridized at 37°C for 16 hrs-3 days. After hybridization, cells were washed 3 x 5 min in 2x SSC at RT, and for 3 x 5 min in 0.1x SSC at 60°C, and then rinsed with SSCT (4x SSC with 0.2% TWEEN 20) at RT. FISH signals were detected by incubation with Alexa Fluor 647 conjugated Strepavidin (1:200; Jackson ImmunoResearch, Cat. # 016-600-084) or Alexa 594 conjugated Strepavidin (1:200; Life Technology, Cat. # S11227) for biotin-labeled probes, or Alexa Fluor 647 conjugated IgG fraction monoclonal mouse anti-digoxin (1:200; Jackson ImmunoResearch, Cat. # 200-602-156) for digoxigenin labeled probes, diluted in SSCT with 1% Bovine Serum Albumin (MilliporeSigma, Cat. # A7906), for 40 min-2 hrs at RT. Coverslips were washed in SSCT for 4 × 5 min, rinsed with 4x SSC and mounted.

### Mitotic FISH

Metaphase spreads were prepared according to a published protocol (71) with small modifications. Cells grown to 70-80% confluence were incubated with 0.1 μg/ml Colcemid (Thermo Fisher Scientific, Cat. # 15212012) in growth media for ∼1 hr. Cells were then harvested and swollen by incubation in 0.075 M KCl for 10-20 min at 37°C, followed by fixation with freshly prepared Carnoy’s fixative (3:1 v/v ratio of methanol/acetic acid). Chromosomal spreads were made by dropping the fixed swollen cells onto cold wet glass slides. DNA FISH of mitotic spreads was performed using a published protocol (71).

### Microscopy and image analysis

For examining EGFP-LacI signals cells were grown on coverslips and fixed with 3-4% paraformaldehyde in DPBS before mounting. For examining the expression of the three reporter minigenes, SNAP tagged-Lamin B1, SNAP-tagged Fibrillarin and mCherry-Magoh, the cells were first labeled with cell-permeable substrate SNAP-Cell Fluorescein (New England Biolabs, Cat. # S9107S) overnight at 240 nM concentrations. To reduce background of unreacted SNAP-tag substrate, cells were incubated 3x 30 mins with media in the incubator, washed with PBS, and fixed with freshly prepared 4% paraformaldehyde in PBS for 15 min at RT. All samples-including fixed cells expressing fluorescently tagged transgenes, 3D DNA FISH, and mitotic FISH sample-were mounted with a Mowiol-DABCO anti-fade medium (72) containing ∼3 μg/ml DAPI (MilliporeSigma, Cat. # D9542).

3D z-stack images were acquired using a Deltavision wide-field microscope (GE Healthcare), equipped with a Xenon lamp, 60X, 1.4 NA oil immersion objective (Olympus) and CoolSNAP HQ CCD camera (Roper Scientific) or a V4 OMX (GE healthcare) microscope, equipped with a 100X, 1.4 NA oil immersion objective (Olympus) and two Evolve EMCCDs (Photometrics). Images were deconvolved using the deconvolution algorithm (72) provided by the *softWoRx* software (GE Healthcare). Gamma = 0.5 was applied to green channels in Figure 5e, Supplementary Figure S8 and Supplementary Figure S10 for proper display of spots with relatively low signals. All image analysis and preparation were done using Fiji (73). Images were assembled using Illustrator (Adobe), Photoshop (Adobe), or GIMP.

For estimation of episome size, the z-sections containing focused episome images for the DAPI and FISH channels were selected manually from the deconvolved z-stack image. Chromosomes and FISH spots were segmented by applying the k-mean clustering algorithm (number of clusters = 3, cluster center tolerance = 0.0001, randomization seed = 48) from the IJ Plugins Toolkit (http://ij-plugins.sourceforge.net/plugins/toolkit.html). The smallest chromosome was identified by manually searching for the chromosome with the smallest area. Segmented FISH spots overlapping or touching chromosomes were removed manually. Integrated DAPI intensities of the smallest chromosome and of the FISH spots not overlapping or touching chromosomes were calculated by Equation 8 (Mean gray value and Area were measured by Fiji). Average episome size was calculated by Equation 9 (n is the number of FISH spots, chro is the smallest chromosome found in the field, 61.4 Mb is the size of chr19 in mm10).

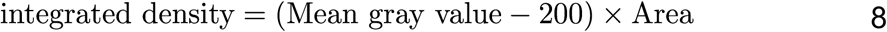

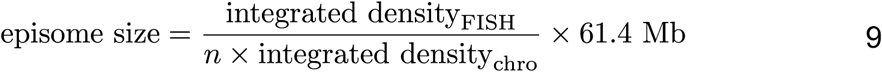

Comparison of reporter gene expression levels in for NIH 3T3 cell clones (Figure 7c) was done by projecting deconvolved images stacks and then measuring the integrated intensity within individual nuclei after subtracting background intensity levels measured in the cytoplasm. Regions of interest circumscribing individual nuclei were drawn manually based on the SNAP-lamin B1 signal. Linear correlations of the integrated intensities of the nuclear SNAP-tag and mCherry signals were calculated using Microsoft Excel with the y-intercept fixed to 0.

A non-linear Gamma correction (0.7) to reduce the grey-scale dynamic range followed by a maximum intensity projection of 3-4 z-sections was used to better visualize both lamin and nucleolar staining simultaneously (Figure 7d).

### Agarose embedded DNA preparation and S1 Nuclease digestion

Agarose embedded DNA was prepared according to published protocols (74, 75) with modifications. To prepare mammalian cell suspensions, cells were grown without selection for 3-4 days after passaging, reaching 80%-90% confluence. Cells were trypsinized, resuspended in cell media, washed with PBS, and resuspended in PBS at a concentration of ∼8 x 10^6^ cells / 100 μl. To prepare *E. coli* cell suspensions, ∼0.1 ml of overnight culture was diluted in 15 ml fresh LB and grown to an OD_600_ of ∼1. Cells were washed with L Buffer (10 mM Tris-Cl pH7.6, 20 mM NaCl, 100 mM EDTA) once and resuspended in L Buffer at a concentration of ∼10^9^/100 μl, assuming a cell concentration of ∼8 x 10^7^/100 ml at an OD_600_ of 1.

2% certified low melt agarose (Bio-Rad Laboratories, Cat. # 1613111) was prepared with L Buffer and kept at 75°C. Equal volumes of the cell suspension (RT) and the agarose solution (75°C) were mixed and immediately transferred to plug molds (Bio-Rad Laboratories, Cat. # 1703713), ∼100 μl mixture per plug. The agarose plugs were incubated in L Buffer with 1% Sarcosyl (MilliporeSigma, Cat. # L5125) and 0.5 mg/ml proteinase K at 55°C for 1-2 days. The agarose plugs were washed with W Buffer (20 mM Tris-Cl, pH7.6, 50 mM EDTA) for 2 x 15 min, incubated in 1 mM PMSF in W Buffer for 30 min, and washed with W Buffer again. Prepared agarose plugs were stored in 0.5 M EDTA at 4°C before use.

For S1 Nuclease (Promega, Cat. # M5761) digestion, agarose plugs were first washed in TE (10 mM Tris-Cl, 1 mM EDTA, pH 7.6) for 3 x 10 min and in 1x S1 Nuclease Buffer for 20 min on ice. The agarose plugs were then digested with 1-16 U/0.4 ml S1 Nuclease in 1x S1 Nuclease Buffer at 37°C for 45 min. The reaction was stopped by washing the agarose plugs with 0.5 M EDTA or W Buffer.

### Pulsed Field Gel Electrophoresis (PFGE)

PFGE was performed using a CHEF-DR III (Bio-Rad Laboratories) according to the manufacturer’s manual using a 1% certified megabase agarose (Bio-Rad Laboratories, Cat. # 1613108) gel in 0.5x Tris-borate-EDTA buffer (TBE), a 0.5x TBE running buffer, and the following parameters: voltage = 6 V/cm, angle = 120°, pulse = 60-120 sec, temperature = 14 °C, run time = 20 or 24 hrs (stopped at 18-20 hrs). Yeast chromosomes (Bio-Rad Laboratories, Cat. # 170-3605) were used as DNA size markers.

### Southern hybridization probes

Southern hybridization probes were created and labeled with digoxigenin by PCR using primers listed in Supplementary Table S1. Set 1 contains a 620bp and a 615 bp fragment amplified from the GFP-ZeoR region; Set 2 contains 525 bp, 534 bp, and 504bp fragments amplified from the BAC vector region; Set 3 contains 446 bp, 681 bp, and 424 bp fragments amplified from the HBB BAC. Pooled Set 1 and Set 2 fragments were used for detecting the DHFR BAC, and pooled Set 1 and Set 3 for detecting the HBB BAC. PCR was done using *Taq* DNA polymerase (New England Biolabs, Cat. # M0267L) with the following recipe: 1x ThermoPol Buffer, 0.2 mM dATP/dCTP/dGTP (New England Biolabs, Cat. # N0446S), 0.165 mM dTTP (New England Biolabs, Cat. # N0446S), 0.035 mM Digoxigenin-11-dUTP, 0.5 ng HBB BAC, 1.25 U *Taq* DNA polymerase, 0.5 μM forward/reverse primers, 50 μl total reaction volume. PCR products were column (QIAGEN, Cat. # 28104) purified. Pooled probes were denatured in nuclease free water, at ∼100°C for ∼10 min and snap-chilled on ice before use.

### Southern hybridization

Southern blotting used a published protocol (76) with modifications. After ethidium bromide staining and imaging, the gel was depurinated in 0.25 M HCl for 2x 30 min, denatured in 0.4 M NaOH for 2x 25 min, neutralized in 0.5 M Tris-Cl/1.5 M NaCl (pH 7.6) for 2x 20 min and washed in 2x SSC for 2x 20 min. DNA was transferred to Zeta-Probe membranes (Bio-Rad Laboratories, Cat. # 1620165) using a Model 785 Vacuum Blotter (Bio-Rad Laboratories), with 2x SSC as transfer buffer, ∼5 inches Hg pressure, and ∼16 hrs transfer time. A Stratalinker (Strategene) was used to cross-link DNA to the membrane.

Hybridization used a standard protocol (77) with modifications. The hybridization buffer was composed of 1:1 volumes of 1 M Na_2_HPO_4_ (pH 7.2) and 14% (w/v) SDS. Total concentration of pooled probes was ∼100 ng/ml. Hybridization was carried out at 65°C for ∼16 hrs. After hybridization, the membrane was washed with 2x SSC/0.1% SDS for 2 x 5 min at room temperature, and with 1x SSC/0.1% SDS for 2 x 10 min at 65°C and rinsed with 2x SSC. Signals were detected using the DIG Nucleic Acid Detection Kit (MilliporeSigma, Cat. # 000000011175041910) according to the manufacturer’ manual, except that in the final step, CDP-*Star* (MilliporeSigma, Cat. # 11685627001) was used instead of NBT/BCIP, and the membrane was imaged by an iBright system (Thermo Fisher Scientific).

### Estimation of average BAC DNA content per episome

To estimate the average BAC DNA content per episome of clone DHFR-UG-s3 and clone HBB-UG-100d3, cells at the same passage were seeded on glass coverslips for DNA FISH using BAC probes, and in different plates for genomic DNA extraction followed by qPCR. The mean number of FISH spots per nucleus, counted from z-stack projected images, provided the average episome copy number per cell. For the DHFR-UG clone, 3 was subtracted from the mean number of FISH spots, as the parental NIH 3T3 cells had ∼3 FISH spots, corresponding to the endogenous DHFR loci, using FISH probes prepared from the DHFR BAC. qPCR estimation of BAC copy number per cell was described in section “Estimation of transgene copy number”. BAC DNA content per episome was calculated using equation 10.

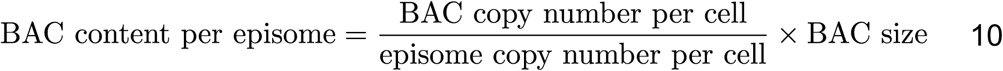

### Whole genome sequencing

Clone DHFR-UG-s3 and clone HBB-UG-100d3 were sorted by flow cytometry using the H1, H2 and L sorting windows shown in Figure 6c and Supplementary Figure S4a. Cells from the H2 and L regions were sorted in the same experiment, while cells from the H1 regions were sorted in another experiment. 100-200 thousand cells were collected from each window. Genomic DNA from sorted cells was isolated by phenol-chloroform extraction. To prepare sequencing libraries, genomic DNA was first fragmented to 100-500 bp by sonication using a Bioruptor Pico (Diagenode), with the following conditions: 4 ng/μl DNA in 120 μl EB, 1.5 ml tube, 10-11 cycles of 30 secs on and 30 secs off. Next, indexed adaptors was attached to the fragmented DNA using True-Seq ChIP Sample Preparation kit (Illumina, Cat. # IP-202-1012) according to the manufacturer’s instructions with the following modifications: after the fragmented DNA was end repaired, 3’ end adenylated, and ligated to indexed adaptors without size selection, the ligation products were PCR amplified for 7∼9 cycles. Libraries were quality checked on a Fragment Analyzer (Agilent) and quantitated by qPCR. Every 6 libraries were pooled at equal molar ratios and sequenced on one lane using a HiSeq 4000 for 101 cycles from one end of the fragments using a HiSeq 4000 sequencing kit version 1. Fastq files were generated and de-multiplexed with the bcl2fastq v2.20 Conversion Software (Illumina). Library quality checking, quantitation and sequencing, and fastq file generation and de-multiplexing were done by the DNA services lab, Roy J. Carver Biotechnology Center, UIUC. 59-65 million reads with quality score >30 were obtained for each library.

### Sequencing reads processing and copy number variation analysis

Low quality bases and adaptor sequences were trimmed from raw reads using cutadapt 1.14 with Python 2.7.13 with the following parameters: -a AGATCGGAAGAGCACACGTCTGAACTCCAGTCAC -q 20,20 -m 20, resulting in ∼0.2% bp being trimmed. Reads were then aligned to a reference genome (mm10 plus HBB BAC (CTD-2643I7, sequence from hg38), the BAC vector (pBelo11, GenBank Accession #: U51113) and UBC-GFP-ZeoR, each as an individual chromosome) using Bowtie2 (version 2.3.2) with default parameters. Overall alignment rate of each sample was ∼98%-99%. Finally, PCR duplicates were removed by SAMtools rmdup (version 1.7) with default parameters, resulting in 42-48 million total mapped reads in each sample.

For reads binning, each chromosome of the reference genome was divided into non-overlapping 3 kb or 30 kb bins; the number of alignments with centers falling into each bin (binned reads) was counted and then divided by the mean read count (Equation 11), generating normalized binned reads (normalized reads, Equation 12), and finally the normalized binned reads of the test sample (H1 or H2 cells) were divided by that of the reference sample (L cells), generating the ratio of normalized binned reads (ratio, Equation 13). The mean read count was ∼50 or ∼500 for 3 kb or 30 kb bin size, respectively. To reduce noise caused by extremely low read counts, a threshold for determining outliers was calculated based on the quantile range (Equation 14). Bins with log_2_(reads) smaller than the threshold in the test sample were removed from further analysis. The excluded bins took up ∼6.0% of total bins for both 3 kb and 30 kb bin sizes, including zero read count bins, which took up ∼5.5% or ∼3.5% of total bins for 3 kb or 30 kb bin size, respectively. The maximum number of reads of the excluded bins were ∼7 or ∼108 for 3 kb or 30 kb bin size, respectively.

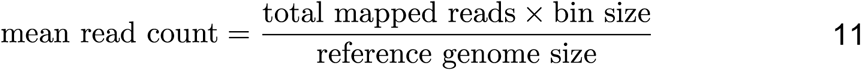

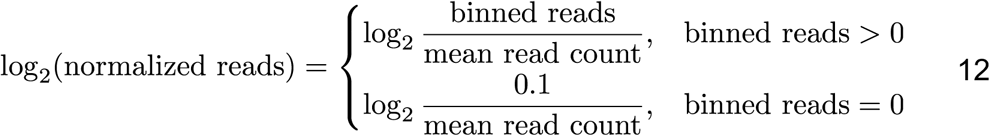

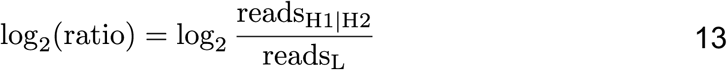

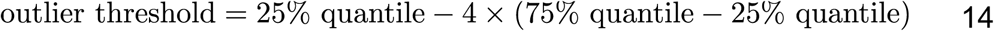

A circular binary segmentation algorithm (78, 79) from the R-package DNAcopy (version 1.52.0) was used to merge bins with similar log_2_(ratio) into segments, with the following parameters for the segment function: verbose = 1, undo.splits=“sdundo”, undo.SD=1. The mean log_2_(ratio) of each segment was calculated for identifying episome-localizing regions.

To identify possible episome-localizing regions, we first measured BAC transgene copy numbers in the H1, H2 and L samples by qPCR and then calculated the theoretical episome copy numbers using the estimated BAC copy number per episome of unsorted cells (Table 2). The minimum copy number increase of episome-localizing host DNA (minimum increase) was then calculated assuming NIH 3T3 to be tetraploid and each episome to have the same host DNA sequence (Equation 15). Segments with mean log_2_(ratio) equal to or greater than log_2_(minimum increase) in both H1 and H2 samples were selected as candidate episome-localizing regions.

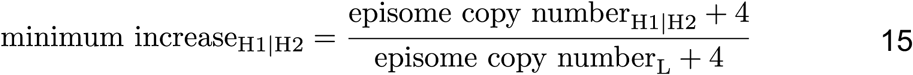

### Construction of DHFR BAC deletions

We tested several DHFR BAC deletions-made for other purposes- for their ability to produce episomes. The DHFR-c27 BAC (51) containing a 256-mer Lac operator (LacO) repeats and a CMV-mRFP-SV40-ZeoR expression cassette was derived from the DHFR BAC, and was used for making the DHFR BAC deletions. DHFR-c27d2 contains a ∼70 kb deletion of the 3’ part of the *Msh3* gene. DHFR-c27d3-crz contains a ∼80 kb deletion of the whole *Dhfr* gene and the 5’ part of the *Msh3* gene, including the CMV-mRFP-SV40-ZeoR expression cassette inserted in the *Msh3* gene, and contains a new CMV-mRFP-SV40-ZeoR expression cassette introduced at the remaining part of *Msh3* gene. DHFR-c27d4 contains a ∼20 kb deletion around the divergent promoter region. *λ* Red-mediated BAC recombineering with a *galK*-based dual-selection scheme was used to create the deletion BACs from the DHFR-c27 BAC, as described in “Construction of dual reporter DHFR BACs” and “Construction of BACs containing the UBC-GFP-ZeoR cassette”. DNA fragments containing either GalK or FRT-GalK-FRT and homology ends were produced by PCR using either pGalK or pUGG as templates. For DHFR-c27d2, the GalK cassette was introduced by the first round of recombination and was subsequently removed by another round of recombination using a DNA fragment created by a pair of partially overlapping primers. For DHFR-c27d3 and DHFR-c27d4, the FRT-GalK-FRT cassette was introduced in the first round instead and was subsequently removed by inducing FLP recombinase as described in “Construction of BACs containing the UBC-GFP-ZeoR cassette”. To create DHFR-c27d3crz, the CMV-mRFP-SV40-ZeoR cassette was introduced into DHFR-c27d3 by one round of recombination using Zeocin resistance as positive selection as described in “Construction of dual reporter DHFR BACs”. Each round of recombination was validated by PCR and restriction enzyme fingerprinting. All primers are listed in Supplementary Table S1.

### Construction of multi-reporter DHFR BAC by BAC-MAGIC

#### Overview

Construction of the 3-reporter BAC was done by serially inserting ∼10-15 kb DNA cassettes into the DHFR BAC scaffold by BAC recombineering. These DNA cassettes were constructed from two different DNA plasmid module types: reporter modules and intervening DHFR sequence modules. DNA cassettes were inserted sequentially into the DHFR BAC using multiple rounds of BAC recombineering and positive selection with one of two different positive selectable markers. After insertion of the first DNA cassette, each subsequent insertion of the next DNA cassette removed the preceding positive selectable located at the 3’ end of the preceding cassette while inserting the alternative selectable marker located at the 3’ end of the new cassette. Three reporter gene modules (Rep Mod 01, 02, 03) plus three intervening DHFR sequence modules (DHFR 02, 03, 04) were constructed and then inserted into the DHFR BAC using 6 sequential rounds of BAC recombineering. In this way, 45 kb of the original DHFR BAC effectively was reconstructed such that the original DHFR sequences were retained but the 3 reporter mini-genes were inserted into this BAC region with each reporter minigene spaced by ∼10 kb of DHFR sequence. We call this overall construction approach BAC-MAGIC (**<u>BAC</u>**-**<u>M</u>**odular **<u>A</u>**ssembly of **<u>G</u>**enomic loci **I**nterspersed **<u>C</u>**assettes).

Each DNA cassette was constructed using traditional cloning methods, Gibson assembly (80), and/or DNA Assembler (81, 82). Three reporter recipient modules (pRM01-Spec, pRM02-Spec, and pRM03-Spec) were designed to incorporate a rare AgeI restriction site for insertion of reporter expression cassettes of choice, in order to create the final reporter modules for BAC recombineering. Unless mentioned specifically all the enzymes were procured from New England Biolabs. All primers and oligos are listed in Supplementary Table S1. Gibson assembly used Gibson assembly cloning kit (New England Biolabs, Cat. # E5510S) as per the manufacturer’s instructions.

DNA Assembler used *Saccharomyces cerevisiae (S. cerevisiae)* strain VL6-48N (MATα, his3-Δ200, trp1-Δ1, ura3-Δ1, lys2, ade2-101, met14, cir°), transformed with 43 fmol pRS413 vector backbone and 130 fmol of all other fragments using the LiAc/SS carrier DNA/PEG method (83). The *S. cerevisiae* single-copy shuttle vector pRS413 contains CEN6/ARS autonomously replicating sequence, auxotrophic selection marker *HIS3* for propagation in yeast, and pMB1 origin of replication and *bla* (Ap^R^) marker for selection with ampicillin in *E. coli*. The 3.8 kb pRS413 vector backbone was PCR amplified from plasmid pRS413 (New England Biolabs) using primer pair RS413-Fw/RS413-Rev for all yeast assembly reactions. The vector backbone and all other fragments made by PCR were digested with DpnI to remove template DNA. Transformants were selected on SC selection media plates lacking histidine [0.17% Bacto-yeast nitrogen base without amino acids (MilliporeSigma, Cat. # Y1251-100G), 0.5% ammonium sulfate, 2% D-glucose, 0.2% Dropout mix (MilliporeSigma, Cat. # Y2001-20G), 2% agar, 80 mg/l uracil, 80 mg/l L-tryptophan, and 240 mg/l L-leucine] at 30°C for 3-4 days. Plasmid DNA were prepared using QIAprep Spin Miniprep Kit (Qiagen, Cat. # 27104) and screened by restriction enzyme fingerprinting. Plasmid DNA from selected yeast colonies was introduced into *E. coli* strain DH5*α* and isolated plasmid DNA then further validated by additional restriction enzyme fingerprinting.

Below we describe construction of each reporter and intervening spacer modules and BAC recombineering assembly of these modules to create the 3-reporter BAC. ApE (M. Wayne Davis, University of Utah, http://biologylabs.utah.edu/jorgensen/wayned/ape/) and SnapGene (from GSL Biotech; available at snapgene.com) programs were used to analyze sequence data, design primers, and design cloning strategies.

#### Construction of plasmid pRM01-RSLB1-Spec (Reporter module 01)

Plasmid pRM01 was made by sequential addition of two DHFR homology regions to plasmid pEGFP-C1 (Clontech). First, the 2.1 kb DHFR homology region (M1F4) was PCR amplified from the DHFR BAC using primer pair M1F4-BamHIfor/M1F4-AgeIrev, double digested with BamHI/AgeI, and ligated with the BamHI/AgeI digested pEGFP-C1 to generate intermediate plasmid pEG-Rep-Module-1a. Next, the 2.0 kb DHFR homology region (M2F12) was PCR amplified from the DHFR BAC using primer pair M2F12-AgeIFor/M2F12-PshRev, double digested with AgeI/PshAI and ligated with the AgeI/SnaBI digested plasmid pEG-Rep-Module-1a to produce plasmid pRM01.

To create plasmid pRM01-Spec (Reporter recipient module 01), a 1.6 kb Spectinomycin resistance gene expression cassette (SpecR), derived from plasmid pYES1L (Thermo Fisher Scientific), was inserted into pRM01, 400 bp upstream of the 3’ end of the M2F12 DHFR homology region by two-fragment Gibson Assembly (80). The two fragments for Gibson assembly were PCR amplified from pRM01 using primer pair GA-RM01-Spec-For/ GA-RM01-Spec-Rev (PCR product size: 7.8 kb), or from pYES1L using primer pair Specfor/SpecRev (PCR product size: 1.6 kb) respectively.

The pRSLB1 (hRPL32-SNAP-Lamin B1) plasmid harboring SNAP-tagged Lamin B1 reporter expression cassette (RSLB1) was constructed by three-fragment Gibson assembly. pEGFP-Lamin B1 plasmid vector backbone 5.3 kb fragment was prepared by AseI/BsrGI double digestion. The hRPL32 promoter (2.2 kb) and SNAP tag (561 bp) fragments were PCR amplified using primer pairs GA-hRPL32-fwd/GA-hRPL32-rev (template plasmid pMOD-HB2-hRPL32-RZ, made in this study), and GA-SNAP-fwd/GA-SNAP-rev (template plasmid pSNAPf, New England Biolabs).

pRM01-Spec was linearized by AgeI and simultaneously dephosphorylated by Shrimp Alkaline Phosphatase (New England Biolabs, Cat. # M0371S). The RSLB1 expression cassette was PCR amplified from plasmid pRSLB1 using primer pair R32CerLBAgeIfor/newPCFAgeIrev (PCR product size: 4.9 kb) and double digested with DpnI/AgeI. The linearized pRM01-Spec and the digested RSLB1 PCR product were ligated to produce plasmid pRM01-RSLB1-Spec, which was digested with AseI to produce the final BAC recombineering 10.3 kb targeting construct.

#### Construction of plasmid pRM02-PSF-Spec (Reporter module 02)

Plasmid pRM02 was made using similar cloning steps used to produce pRM01 except two different DHFR homology regions were added to pEGFP-C1: 2.0 kb PCR product M2F4 (primer pair M2F4-BamHIfor/M2F4-AgeIrev) replaced M1F4 and 2.0 kb PCR product M3F1 (primer pair M3F1-AgeIFor/M3F1-PshRev) replaced M2F12. Plasmid pRM02-Spec was made the same way as pRM01-Spec except that fragment 1 for Gibson assembly was PCR amplified from plasmid pRM02 using primer pair GA-RM02-Spec-For/GA-RM02-Spec-Rev (PCR product size: 7.8 kb). The final plasmid pRM02-Spec (pRep-module 02-Spec) is Reporter recipient module 02 for the SNAP-tagged Fibrillarin reporter expression cassette (PSF).

To create plasmid pPSF (pPPIA-SNAP-Fibrillarin), the GFP cassette between KpnI/HpaI restriction sites of plasmid GFP-Fibrillarin was replaced with a 730 bp Cerulean cassette PCR amplified from plasmid pCerulean-N1 (New England Biolabs) using primer pair ForCerFib/RevCerFib, resulting in an intermediate plasmid pPCF. Next, the CMV promoter between SnaBI/HindIII sites of pPCF was replaced with the 2.8 kb PPIA promoter PCR amplified from plasmid p[MOD-HB2-PPIA-RZ] (made in this study) using primer pair PPIACerFibFor/ PPIACerFibRev, resulting in plasmid pPPIA-Cer-Fib. Finally, the 720 bp Cerulean cassette between the AgeI/HpaI sites of pPPIA-Cer-Fib was replaced with a 560 bp SNAP tag fragment PCR amplified from plasmid pSNAPf (New England Biolabs) using primer pair Snap-XmaI-For/Snap-HpaI-Fib-Rev and double digested with XmaI/HpaI, producing pPSF.

pRM02-Spec was linearized by AgeI and simultaneously dephosphorylated by Shrimp Alkaline Phosphatase (New England Biolabs, Cat. # M0371S). The 4.6 kb PSF expression cassette was PCR amplified from pPSF using primer pair PSF-AgeI-For/ PSF-AgeI-Rev and double digested with DpnI/AgeI. Their ligation produced plasmid pRM02-PSF-Spec, which provided the 10.4 kb BAC recombineering targeting construct after BamHI/AatII/RsrII triple digestion of pRM02-PSF-Spec.

#### Construction of plasmid pRM03-PCM-Spec (Reporter module 03)

Plasmid pRM03 was made using similar cloning steps used to produce pRM01 except two different DHFR homology regions were added to pEGFP-C1: 2.1 kb PCR fragment M3F4 (primer pair M3F4-BamHIfor/M3F4-AgeIrev) replaced M1F4 and 2.1 PCR fragment M4F1 (primer pair M4F1-AgeIFor/M4F1-PshRev) replaced M2F12. Plasmid pRM03-Spec was made the same way as pRM01-Spec except that fragment 1 for Gibson assembly was PCR amplified from plasmid pRM03 using using primer pair GA-RM03-Spec-For/ GA-RM03-Spec-Rev (PCR product size: 7.8 kb). The final plasmid pRM03-Spec (pRep-module 03-Spec) is Reporter recipient module 03 for the mCherry-tagged Magoh reporter expression cassette (PCM).

Plasmid pPCM (pPPIA-mCherry-Magoh) was created in two steps. First, the CMV promoter between the NdeI/NheI sites of plasmid pmRFP-Magoh was replaced with the PPIA promoter (2.8 kb), PCR amplified from plasmid pMOD-HB2-PPIA-RZ using primer pair PPIA-Magohfor/ PPIA-MagohRev and double digested with NdeI/NheI, resulting in intermediate plasmid pPMM. Next, the mRFP tag between the NheI/HindIII sites of pPMM was replaced with a 720 bp mCherry tag PCR amplified from plasmid pQCXIN-TetR-mCherry using primer pair mCherry-NheI-Magoh-For/mCherry-H3-Magoh-Rev, resulting in plasmid pPCM.

To create plasmid pRM03-PCM-Spec (Reporter module 03), plasmid pRM03-Spec was linearized by AgeI and simultaneously dephosphorylated by Shrimp Alkaline Phosphatase (New England Biolabs, Cat. # M0371S). A 4.2 kb PCM expression cassette was PCR amplified form plasmid pPCM using primer pair MMorCF-AgeIfor/newPCFAgeIrev and double digested with DpnI/AgeI. pRM03-Spec and the PCM PCR product were ligated, producing plasmid pRM03-PCM-Spec, which was used as a template for PCR amplification using primer pair M3F4-PCR-Fw/M4F1-PCR-Rev to produce the 9.9 kb BAC recombineering target. After PCR, any remaining template plasmid was digested with DpnI.

#### Construction of plasmid pRS413-DHFR-Mod-02-Kan (Intervening DHFR module 02)

Plasmid pRS413-DHFR-Mod-02 was made by assembling the vector backbone with four additional fragments using the DNA assembler method (81, 82). Fragment 5’-DHM2 (4.3 kb) and fragment 3’-DHM2 (6.3 kb) with an overlap of 659 bp and were both PCR amplified from the DHFR BAC, using primer pair M2F12-AgeIfor/M2F1rev or DHM2-Seq2/M2F4-AgeIrev, respectively. Two bridging oligomers, with a 125 bp homology to the pRS413 vector backbone, and a 125 bp homology to fragment 5’-DHM2 (oligo M2F1-pRS413) or fragment 3’-DHM2 (oligo M2F4-pRS413) were synthesized at Integrated DNA Technologies, Inc. The final Intervening DHFR module 02, plasmid pRS413-DHFR-Mod-02-Kan, was created by ligating a 2.4 kb Kan/NeoR cassette derived from DraI digestion of plasmid pEGFP-C1, with the plasmid pRS413-DHFR-Module-02 linearized by DraIII and blunted by DNA Polymerase I, Large (Klenow) Fragment.

For BAC recombineering an 11.7 kb of targeting construct was amplified from plasmid pRS413-DHFR-Mod-02-Kan using primer pair M2F12-AgeIFor/DH2-4rev and purified by gel extraction after DpnI digestion of the template plasmid.

#### Construction of plasmid pRS413-DHFR-Mod-03-Kan (Intervening DHFR module 03)

Plasmid pRS413-DHFR-Mod-03 was made by assembling the vector backbone with four additional fragments using the yeast DNA assembler method. Fragment 5’-DHM3 (6.5 kb) and fragment 3’-DHM3 (5.0 kb) with an overlap of 1553 bp were both PCR amplified from the DHFR BAC using primer pair M3F1-AgeIFor/M3F3-BamHIrev or M3-F3For/M3F4-AgeIRev, respectively. Two bridging oligomers, with a 125 bp homology to the pRS413 vector backbone, and a 125 bp homology to fragment 5’-DHM3 (oligo M3F1-pRS413) or to fragment 3’-DHM3 (oligo M3F4-pRS413), respectively, were synthesized at Integrated DNA Technologies, Inc. The final Intervening DHFR module 03, plasmid pRS413-DHFR-Mod-03-Kan, was created by ligating a 2.4 kb Kan/NeoR cassette derived from DraI digestion of plasmid pEGFP-C1, with the plasmid pRS413-DHFR-Mod-03 linearized by SmaI.

For BAC recombineering a 12.2 kb targeting construct was amplified from plasmid pRS413-DHFR-Mod-03-Kan using primer pair DH3-1for/DH3-4rev and purified by gel extraction after DpnI digestion of the template plasmid.

#### Construction of plasmid pRS413-DHFR-Mod-04-Zeo (Intervening DHFR module 04)

Plasmid pRS413-DHFR-Mod-04 was made by assembling the vector backbone plus 5 additional fragments using the yeast DNA assembler method (4). Fragment 5’-DHM4 (4.9 kb), fragment Mid-DHM4 (5.2 kb) and fragment 3’-DHM4 (5.2 kb) with an overlap of 2663 bp in between 5’-DHM4 and Mid-DHM4, and an overlap of 2542 bp in between Mid-DHM4 and 3’-DHM4, were PCR amplified from the DHFR BAC using primer pair M4F1-AgeIfor/DHM4F2-R, DHM4F2-Fw/DHM4F3-R, or Fw-M4F2-BamHI/RevM4F5-MluI, respectively. Two bridging oligomers, with a 125 bp homology to pRS413 vector backbone, and a 125 bp homology to fragment 5’-DHM4 (oligo M4F1-pRS413), or to fragment 3’-DHM4 (oligo M4F5-pRS413), were synthesized at Integrated DNA Technologies, Inc. The final Intervening DHFR module 04, plasmid pRS413-DHFR-Mod-04-Zeo, was created by ligating a 1.1 kb ZeoR expression cassette PCR amplified from plasmid pSV40/Zeo2 (ThermoFisher Scientific) using 5’ phosphorylated primer pair ZeoMluIFor/ZeoMluIRev, with the plasmid pRS413-DHFR-Module-04 linearized by BmgBI.

For BAC recombineering an 11.6 kb targeting construct was excised out from plasmid pRS413-DHFR-Mod-04-Zeo using KpnI/DrdI restriction enzymes and gel purified.

#### Assembly of modules to create multi-reporter DHFR BAC

The six targeting constructs derived from the three reporter modules and the three intervening DHFR modules were incorporated into the DHFR BAC by BAC recombineering, with the following order: Reporter module 01, Intervening DHFR module 02, Reporter module 02, Intervening DHFR module 03, Reporter module 03 and Intervening DHFR module 04. *E. coli* strain SW102 was used for BAC recombineering. Each round of BAC recombineering used a corresponding antibiotic (50 μg/ml Kanamycin, 50 μg/ml Spectinomycin, or 25 μg/ml Zeocin) as positive selection for incorporation of the current targeting construct as described in section “Construction of dual reporter DHFR BACs”. In the second to the last round of BAC recombineering, colonies were further screened for loss of the antibiotic resistance gene incorporated in the previous round of BAC recombineering by streaking colonies onto a plate containing the corresponding antibiotic. Each round of recombination was validated by restriction enzyme fingerprinting.

## RESULTS

### Overview of BAC TG-EMBED toolkit development

We previously demonstrated the feasibility of the BAC TG-EMBED approach using both the DHFR BAC (51) and a BAC containing the human GAPDH gene locus (GAPDH BAC) (54). We set out to extend this BAC TG-EMBED methodology in two new directions (Figure 1).

**Figure 1.**
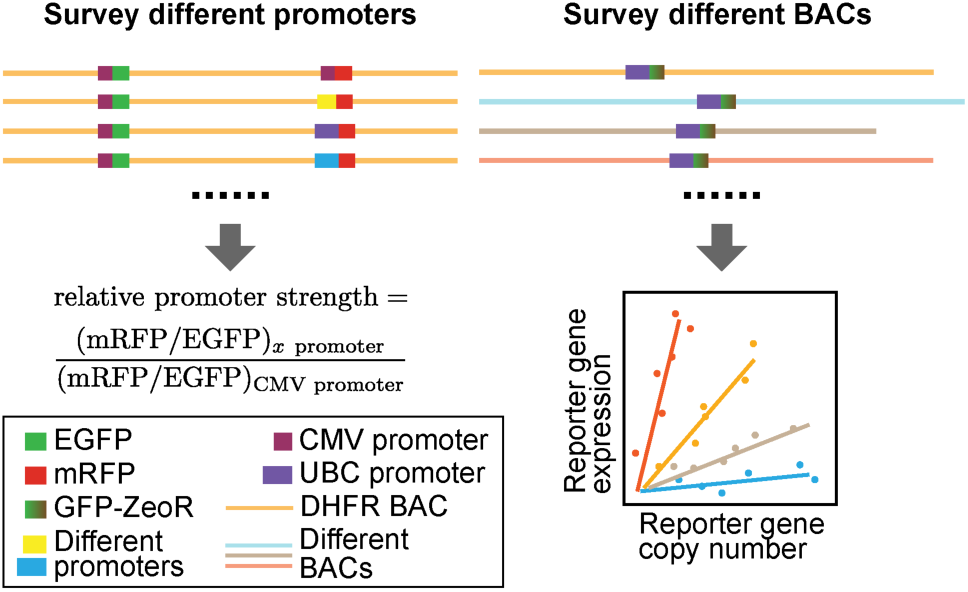
Two-prong experimental approach. Left: Identification of promoters of different strengths-We measured relative promoter strengths by embedding EGFP and mRFP reporter genes into the DHFR BAC, using the CMV promoter to drive EGFP expression and the test promoter to drive mRFP. The ratio of mRFP and GFP expression, normalized by this same ratio for a CMV test promoter, defines promoter strength relative to CMV. Right: Surveying reporter gene expression in different BAC scaffolds-(Top) The UBC-GFP-ZeoR reporter gene was inserted into BACs carrying DNA from mouse or human genomic regions corresponding to either transcriptionally active or inactive genomic regions. (Bottom) Plotting reporter gene expression (y-axis) versus reporter gene copy number (x-axis) for multiple cell clones stably expressing BAC transgenes: a linear correlation would indicate copy-number dependent, position independent expression, while the slope of this linear correlation would measure reporter gene expression per copy number.

First, to better control transgene expression and to be able to express multiple transgenes at reproducible expression ratios, we explored a set of constitutive promoters with various strengths for transgene expression. A previous similar survey of promoters within BAC scaffolds focused only on strong promoters (53). Moreover this survey compared average expression in pools of cell colonies containing different copy-number BAC insertions (53). Here we used a two-reporter, single-cell ratio assay and also examined promoters with a wide range of promoter strengths. Testing each promoter with each BAC scaffold would have generated too large a number of possible combinations. We therefore decided to test a number of different promoters with the original DHFR BAC.

Second, we used one specific reporter gene construct to survey the effect of different BAC scaffolds on reporter gene expression. Previous similar applications used BAC scaffolds containing multiple endogenous genes which would also be expressed in addition to added transgenes (51–53). Moreover, in a previous, similar application, different strong promoters were tested by insertion into the exon of an active BAC gene (53). Here we compared BAC scaffolds containing expressed genes with BAC scaffolds from gene deserts or regions containing silenced genes. We assayed the level, stability, and reproducibility of the embedded reporter gene expression when inserted into different BAC scaffolds to identify optimal BAC scaffolds for the BAC TG-EMBED system.

### A toolset of 7 endogenous promoters for tuning relative transgene expression levels

We selected 7 endogenous promoters to test, either because of their known ability and use to drive transgene expression in a range of cell types (EEF1α, UBC) (60, 84–86), or because these promoters were from housekeeping genes (RPL32, PPIA, B2M, RPS3A, GUSB) known to be expressed uniformly across a wide range of tissue types (87–90). We amplified 1-3 kb of regulatory regions upstream of the transcription start sites of these genes using either human genomic DNA as a template or, for the UBC promoter, using the pUGG plasmid (54).

To assay relative promoter strength, we used the two-minigene reporter system developed in our previous study in which we compared expression of CMV-driven EGFP and mRFP minigenes inserted in the same mouse DHFR BAC scaffold (51). We previously showed that the mRFP minigene reporter expression varied less than or equal to 2.4-fold when the mRFP reporter was inserted at 6 different positions ranging 3-80 kb away from the EGFP reporter gene location on the same BAC (51). To compare relative promoter strengths, we fixed the insertion positions of mRFP and EGFP, and measured the relative fluorescence levels of mRFP and EGFP when they were both driven by the CMV promoter versus when the mRFP reporter was driven by an endogenous promoter (Figure 1). Thus our assay measured the strength of different endogenous promoters relative to the viral CMV promoter, while also measuring the variation in this relative strength in different cells of a mixed clonal population.

For this assay, the EGFP reporter minigene was inserted 26kb downstream of the Msh3 transcription start site (51) (Figure 2a). PCR-amplified promoters from 7 different housekeeping genes were cloned upstream of the mRFP expression cassette (Figure 2b), and then this mRFP expression cassette was introduced 121 kb downstream of the Msh3 transcription start site by BAC recombineering (Figure 2a), generating the dual reporter DHFR BAC. As a control, we used the dual reporter BAC previously constructed (51) in which the same mRFP cassette driven by the CMV promoter was inserted at this same location 121 kb downstream of the Msh3 start site.

**Figure 2.**
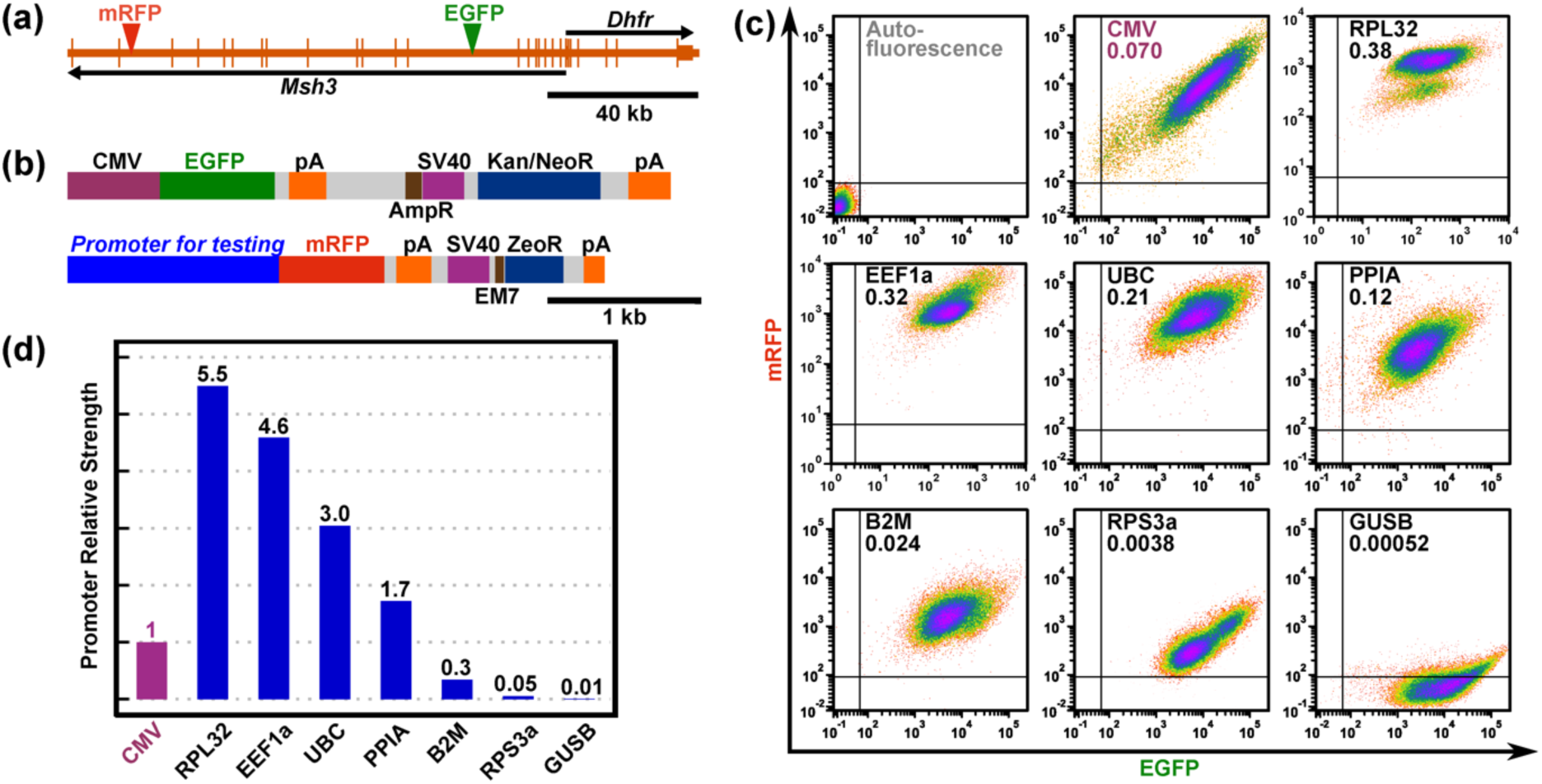
Dual-reporter assay for promoter strength estimation. (a) Dual reporter DHFR BAC showing the two genes on the BAC, *Dhfr* and *Msh3*, and the insertion sites of the two reporter expression cassettes. Longer vertical bars-exons; shorter vertical bars-UTRs; arrows-direction of transcription; green arrowhead-EGFP expression cassette insertion site; red arrowhead-mRFP expression cassette insertion site. (b) The two reporter gene/selectable marker cassettes used in the assay. The EGFP cassette (top) contains an EGFP minigene, driven by a CMV promoter, and a Kanamycin/Neomycin resistance gene (Kan/NeoR), driven by a SV40 promoter for expression in mammalian cells, or by a AmpR promoter for expression in bacteria. The mRFP cassette (bottom) contains a mRFP minigene and a Zeocin resistance gene (ZeoR). Different endogenous promoters were inserted immediately upstream of mRFP. ZeoR is driven by a SV40 promoter for expression in mammalian cells, or by a AmpR promoter for expression in bacteria. pA-poly(A) signal. (c) Scatter plots showing mRFP fluorescence (y-axis) vs EGFP fluorescence (x-axis) of cells from the mixed clonal populations stably transfected with dual reporter DHFR BACs. Promoters driving the mRFP and the ratio of mRFP/EGFP (promoter strength) are labeled in each plot. (d) Promoter strengths relative to CMV.

Mouse NIH 3T3 fibroblasts were then stably transfected with these modified BAC constructs. After dual selection with G418 and Zeocin for two weeks, mixed populations of stable clones carrying the BAC transgenes were analyzed by flow cytometry to measure the relative expression ratio of mRFP and EGFP (Figure 2c). Fluorescent beads were used as an invariant fluorescence standard to calibrate the flow cytometer intensity outputs. The ratio of mRFP to EGFP expression was then normalized by the ratio observed with the original dual-reporter BAC construct in which both reporters were driven by CMV promoter, providing the endogenous promoter strength relative to the CMV promoter.

We observed an overall variation in promoter strength of over 500-fold, ranging from the 4-5 fold relative promoter strength of the RPL32 and EEF1α promoters to the 0.01-fold relative promoter strength for the GUSB promoter as compared to the CMV promoter (Figure 2d). This expression ratio appeared to be similar across the cell population.

### Reporter gene expression as a function of transcriptionally active and inactive BAC scaffolds

To find the best BAC scaffold for the BAC TG-EMBED system, we tested BAC scaffolds from both actively transcribed regions and regions containing silenced genes or no genes. Specifically, we measured the expression as a function of copy number of one specific reporter gene construct inserted into these BAC scaffolds. Previous applications of BAC TG-EMBED showed a linear relationship between copy number and expression level, largely independent of the chromosome integration site, demonstrating copy-number dependent, position independent transgene expression (51, 54). For active chromosomal regions, we chose the RP11-138I1 BAC containing the human ubiquitin B gene locus (UBB BAC), the RP23-401D9 BAC containing the “safe-haven” mouse *Rosa26* genetrap locus (ROSA BAC) (91), and the CITB-057L22 BAC carrying the mouse Dhfr gene locus (DHFR BAC). For inactive chromosomal regions, we chose the CTD-2207K13 BAC (2207K13 BAC) that contains no known gene or regulatory element from a gene-desert region from the human genome, and the CTD-2643I7 (HBB BAC) containing the human HBB gene locus and multiple olfactory genes, all of which are transcriptionally silenced in fibroblasts (92).

We selected the UBC promoter for this reporter gene cassette as this promoter had previously been shown to drive high expression across multiple cell types (86); in our dual reporter system the UBC promoter was 2.6-fold stronger than the CMV promoter (Figure 2d). Moreover, to eliminate any possible transcriptional interference from closely spaced reporter and selectable marker minigenes and to minimize any epigenetic silencing arising from DNA methylation of this reporter gene-selectable marker construct, we used a commercially available GFP-ZeoR fusion protein gene construct in which all CpG dinucleotides had been removed and replaced by synonymous codons (Figure 3a).

**Figure 3.**
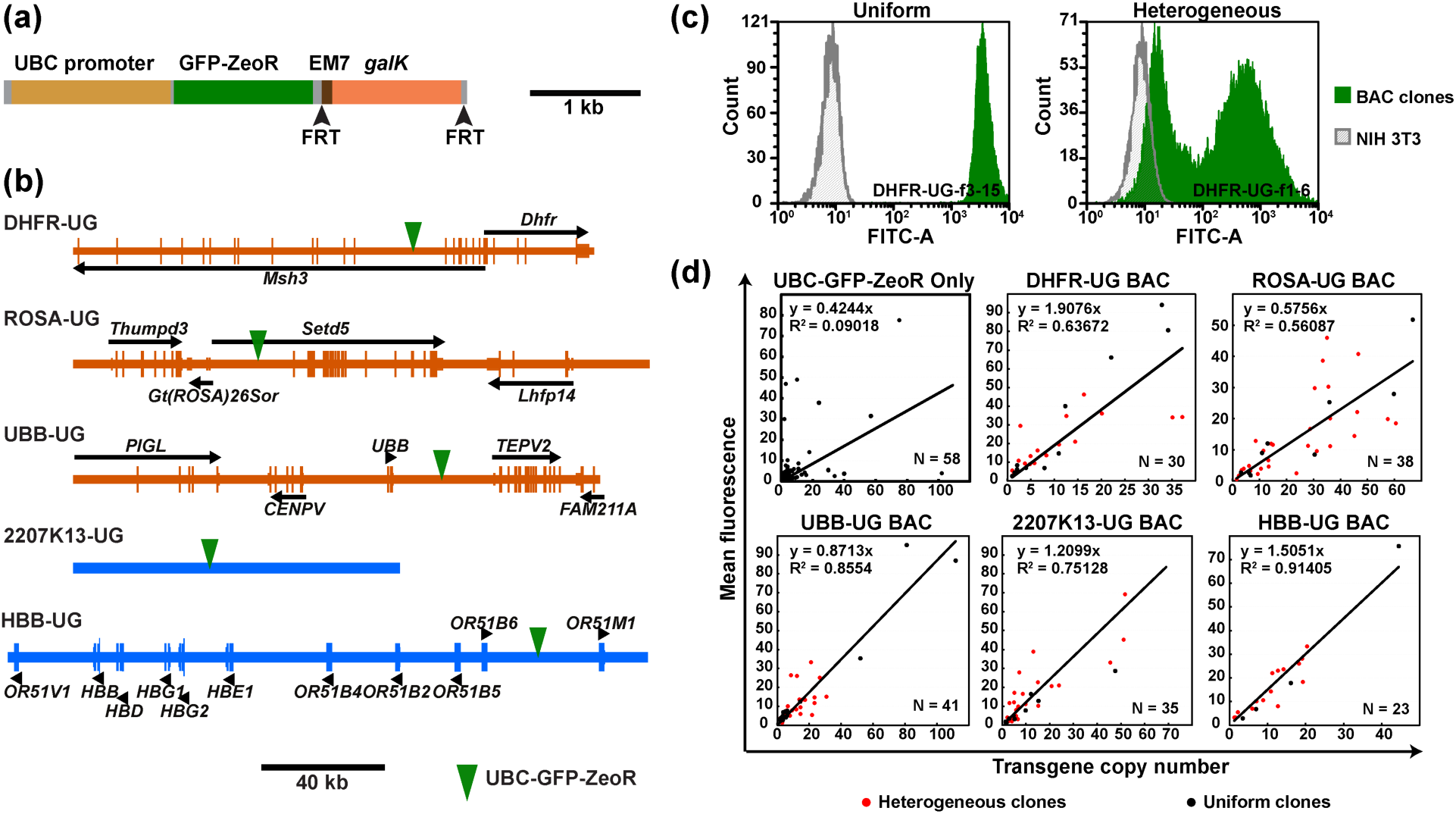
Expression of reporter gene embedded in different BAC scaffolds. (a) UBC-GFP-ZeoR-FRT-GalK-FRT cassette showing the GFP-ZeoR minigene driven by the UBC promoter and the *galK* positive/negative selection marker flanked by 34 bp flippase recognition target (FRT) sites (arrowheads). (b) Maps of the BACs used in the study. Longer vertical bars-exons; shorter vertical bars-UTRs; black arrows or arrowheads-direction of transcription; green arrow heads-UBC-GFP-ZeoR insertion site. (c) GFP fluorescence histograms obtained by flow-cytometry for “uniform” (left, green, clone DHFR-UG-f3-15) versus “heterogeneous” (right, green, clone DHFR-UG-f1-6) expressing NIH 3T3 clones carrying the DHFR-UG BAC. x-axis-fluorescence value, y-axis-cell number; gray-autofluorescence of untransfected cells. Fluorescence is measured in arbitrary units. (d) Scatter plots of mean normalized cellular GFP fluorescence (y-axis) vs reporter gene copy number (x-axis) for clonal populations transfected with the UBC-GFP-ZeoR cassette alone or with different BAC scaffolds carrying the UBC-GFP-ZeoR reporter gene. Linear regression fits (black lines, y-intercepts set to 0) are shown with corresponding R-squared values and equations. Red circles-heterogeneous clones; Black circles-uniform clones; Bottom right of plots: Number of clones analyzed.

We inserted this UBC-GFP-ZeoR reporter gene construct into different BAC scaffolds by BAC recombineering, using *galK* for positive/negative selection (63, 64). To eliminate potential artifacts caused by proximity to active promoters, transcriptional start sites (TSS), or miRNA sequences, we chose insertion sites flanked on both sides by at least 5 kb free of such sequence elements (Figure 3b). The UBB, HBB, 2207K13, ROSA, DHFR BACs with the UBC-GFP-ZeoR reporter gene insertion were named as UBB-UG, HBB-UG, 2207K13-UG, ROSA-UG and DHFR-UG.

After transfection, multiple cell clones (n=20-40) carrying stably integrated BAC arrays were selected for Zeocin resistance and analyzed for reporter gene expression by flow cytometry, using untransfected NIH 3T3 cells to determine background, autofluorescence levels. For each cell clone, we used flow cytometry to measure the mean GFP reporter expression and qPCR to measure reporter gene copy number. These cell clones showed GFP fluorescence mean levels ranging from 10-1000 fold higher than the background autofluorescence.

Our original working hypothesis predicted that the BAC TG-EMBED reporter expression should be uniform in all cells of the same clone. Also, we expected to see a linear relationship between mean reporter gene fluorescence and number of BAC copies, signifying a copy-number-dependent, position independent expression. Furthermore, we expected that the slope of this linear relationship would be higher for BAC scaffolds expected to reconstitute an active chromatin environment permissive for transgene expression as compared to BAC scaffolds expected to reconstitute a more condensed, inactive chromatin environment (Figure 1). In contrast, we expected that the reporter gene cassette transfected without any BAC scaffold would show clonal expression levels that poorly correlated with reporter gene copy number (copy-number-independent expression).

Unexpectedly, the stable cell clones we isolated showed two distinct types of population expression profiles-uniform versus heterogeneous. Uniform clones showed single, relatively narrow expression peaks in the flow cytometry histograms, with more than 90% of the cells showing GFP fluorescence varying only over a 10-fold intensity range (Figure 3c, left). Heterogeneous clones instead showed two peaks with a range of GFP expression varying ∼1000-fold, with the lower GFP intensity peak overlapping with the autofluorescence distribution of control cells (Figure 3c, right). We had not previously observed such heterogeneous expression profile using our original DHFR BAC containing the CMV-driven mRFP alone or both the CMV-driven EGFP and CMV-driven mRFP reporter genes (51). However, we had observed ∼80% uniform clones for a GAPDH BAC scaffold with the UBC-GFP-ZeoR reporter gene inserted (54). The percentage of clones showing such heterogeneous expression varied from 58% to 83% for the 5 BAC scaffolds surveyed here (Table 1). No similar heterogeneous expression profile was observed when the reporter gene construct was transfected by itself (Table 1).

**Table 1.**
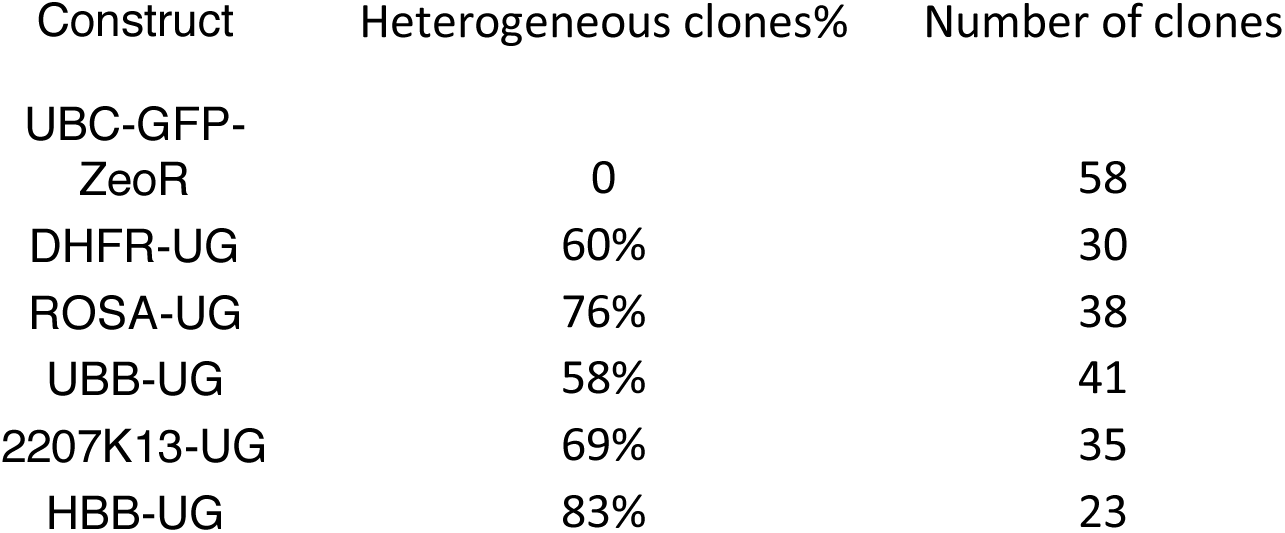
Percentage of heterogeneously expressing clones transfected with the UBC-GFP-ZeoR cassette alone or with different BAC scaffolds carrying the UBC-GFP-ZeoR reporter gene.

As expected, the control transfection of the reporter gene cassette by itself resulted in copy-number-independent expression of the reporter gene (Figure 3d, R^2^=0.09), while the reporter gene embedded within the BACs yielded a linear relationship between reporter gene fluorescence for both uniform (black) and heterogeneous (red) BAC transgene clones (Figure 3d, R^2^=0.561 to 0.914).

Surprisingly, we observed no more than a 4-fold variation in expression per copy number among the 5 different BAC scaffolds tested, with no obvious relationship between the observed slope and the type of BAC scaffold (Figure 3d). Although the transcriptionally active DHFR BAC produced the highest slope, the transcriptionally inactive HBB BAC and the 2207K13 BAC containing DNA from a gene desert produced the second and third highest slopes, while the BAC containing DNA from the “safe haven” mouse *Rosa26* locus produced the lowest slope.

Overall, these results show that for this UBC-GFP-ZeoR reporter gene, high-level, copy-number-dependent transgene expression using the BAC TG-EMBED method does not require BACs containing active, housekeeping genomic regions, but can also be obtained from a wide range of BAC genomic DNA inserts, including gene-desert regions. This means BAC TG-EMBED can be used to drive expression of only the transgenes added to the BAC scaffold, without overexpression of the genes contained within the BAC scaffold.

### Temporal stability of BAC-embedded reporter gene expression in uniform cell clones

We previously showed that the BAC TG-EMBED method provided long-term stability of transgene expression in the presence of continued drug selection (51). However, in the absence of drug selection we observed a 30-80% drop in expression over several months of cell passaging without any apparent drop in the integrated BAC copy number (51).

Here we determined the long-term stability of the UBC-GFP-ZeoR reporter gene expression for both uniform and heterogeneous clones for four different BAC scaffolds. Individual clones for each BAC scaffold (3 uniform and 2 heterogeneous for ROSA-UG BAC, 7 uniform and 4 heterogeneous for 2207K13-UG BAC, 8 uniform and 3 heterogeneous for UBB-UG BAC, and 3 uniform and 3 heterogeneous for DHFR-UG BAC) were passaged up to three months in the absence or presence of drug selection and analyzed for reporter gene fluorescence at regular intervals after removal of drug selection.

With the exception of a small number of apparent fluctuations possibly related to transient changes in culture conditions, clones with uniform reporter gene expression showed no significant change either in the mean fluorescence values (Figure 4a) or in the distribution of fluorescence among the same clones (Figure 4b and Supplementary Figures S1) over time in the absence of selection for all four BAC scaffolds tested. In the presence of continued selection, uniform clones containing DHFR-UG or ROSA-UG BACs showed no significant reporter gene expression change, while an ∼50% or 100% increase was observed for the UBB-UG or 2207K13-UG BAC clones, respectively (Figure 4a). No changes in estimated BAC copy number based on qPCR measurement were observed for any of these clones during this time series. This suggests that epigenetic changes driven by selection pressure may be responsible for these small increases in reporter gene expression.

**Figure 4.**
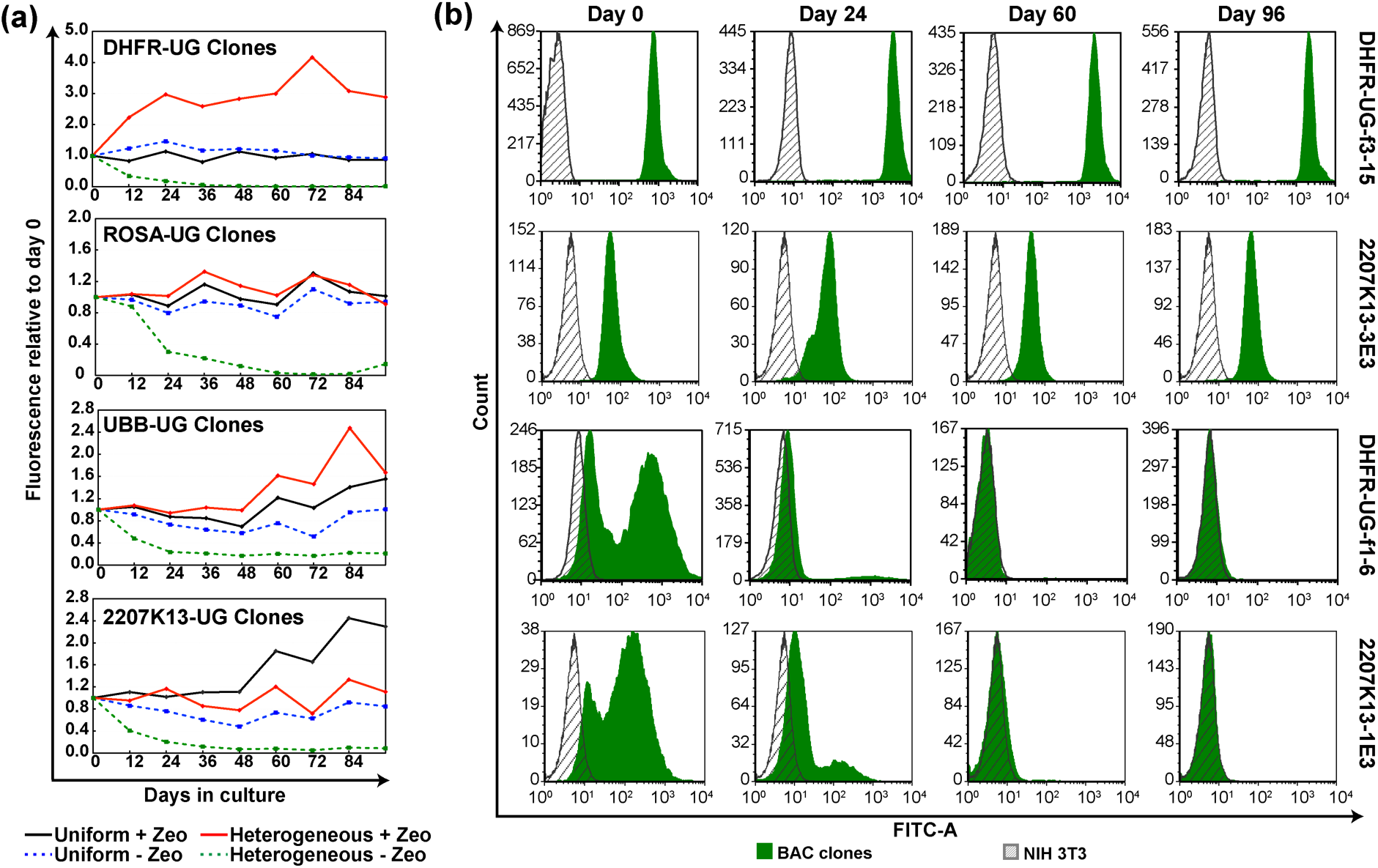
UBC-GFP-ZeoR reporter gene expression over time. “Uniform” clones show stable expression with or without expression, while “heterogenous” clones show progressive loss of expression without selection. (a) Changes in GFP fluorescence of uniform versus heterogeneous clones, averaged over multiple clones (2–8), carrying indicated BAC transgenes during 96 days of continuous passaging with or without Zeocin selection. x-axis-number of days since removal of Zeocin; y-axis-mean fluorescence values of multiple clones divided by that at day zero; black-“uniform” expressing clones cultured with Zeocin; blue-“uniform” expressing clones cultured without Zeocin; red-“heterogeneous” expressing clones cultured with Zeocin; green-“heterogeneous” expressing clones cultured without Zeocin; (b) GFP fluorescence histogram of representative “uniform” and “heterogeneous” expressing NIH 3T3 clones at day 0, 24, 60 and 96 without selection. Gray-autofluorescence of untransfected cells; Green-GFP fluorescence of the indicated clones. x-axis-fluorescence; y-axis-cell number.

Notably, in the absence of selection, heterogeneous clones for all tested BAC scaffolds showed a significant and progressive loss of reporter gene expression over time. This led to a significant fraction of cells showing autofluorescence levels of fluorescence by the end of the experiment (Figure 4a). Reporter gene expression-level became progressively more homogenous, but at lower fluorescence levels (Figure 4b and Supplementary Figure S1). With selection, UBB-UG and DHFR-UG BAC heterogeneous clones showed a 1.6 to 3-fold increase in reporter gene expression, respectively, while the other BAC scaffold heterogeneous clones showed no significant changes (Figure 4a).

### BAC transgenes are maintained as episomes in heterogeneous clones

In our previous work, all stable cell clones obtained after BAC transfection and drug selection contained single BAC copies or multi-copy BAC arrays that had integrated into endogenous chromosomes (51, 65, 93–95) consistent with similar results from numerous laboratories. Thus, we initially assumed that the broad distribution of reporter gene fluorescence observed in heterogeneous cell clones was due to position effect variegation (PEV) of the BAC TG-EMBED reporter genes. We hypothesized that integrations into some chromosome integration sites led to uniformly-expressing clones, while integration into other chromosome sites prone to PEV led to heterogeneous clones with variegated transgene expression.

However, the observation of a progressive loss over time of reporter gene expression for all heterogeneous clones led us to question the genome stability of the BAC transgenes in these clones. To test the relationship between changes in reporter gene expression and BAC copy number, we first sorted cells from the heterogeneous DHFR-UG-s3 cell clone by fluorescence-activated cell sorting (FACS), using a narrow sorting-window centered around the GFP peak fluorescence level (Figure 5a). After cell-sorting, with drug selection the original heterogeneous reporter gene expression distribution reestablished itself within one week of culture (Figure 5b). We then resorted cells showing different levels of GFP fluorescence using four narrow fluorescence windows P1, P2, P3, and P4 (Figure 5b), and then used qPCR to measure the BAC copy number in cells from each of these sorting windows. Plotting mean cell fluorescence intensity levels versus copy number for these clonal subpopulations yielded a strikingly linear relationship (R^2^=0.99) (Figure 5c). Thus, the variable reporter gene expression level in this heterogeneous cell clone is the result of loss of BAC transgenes rather than chromosome PEV.

**Figure 5.**
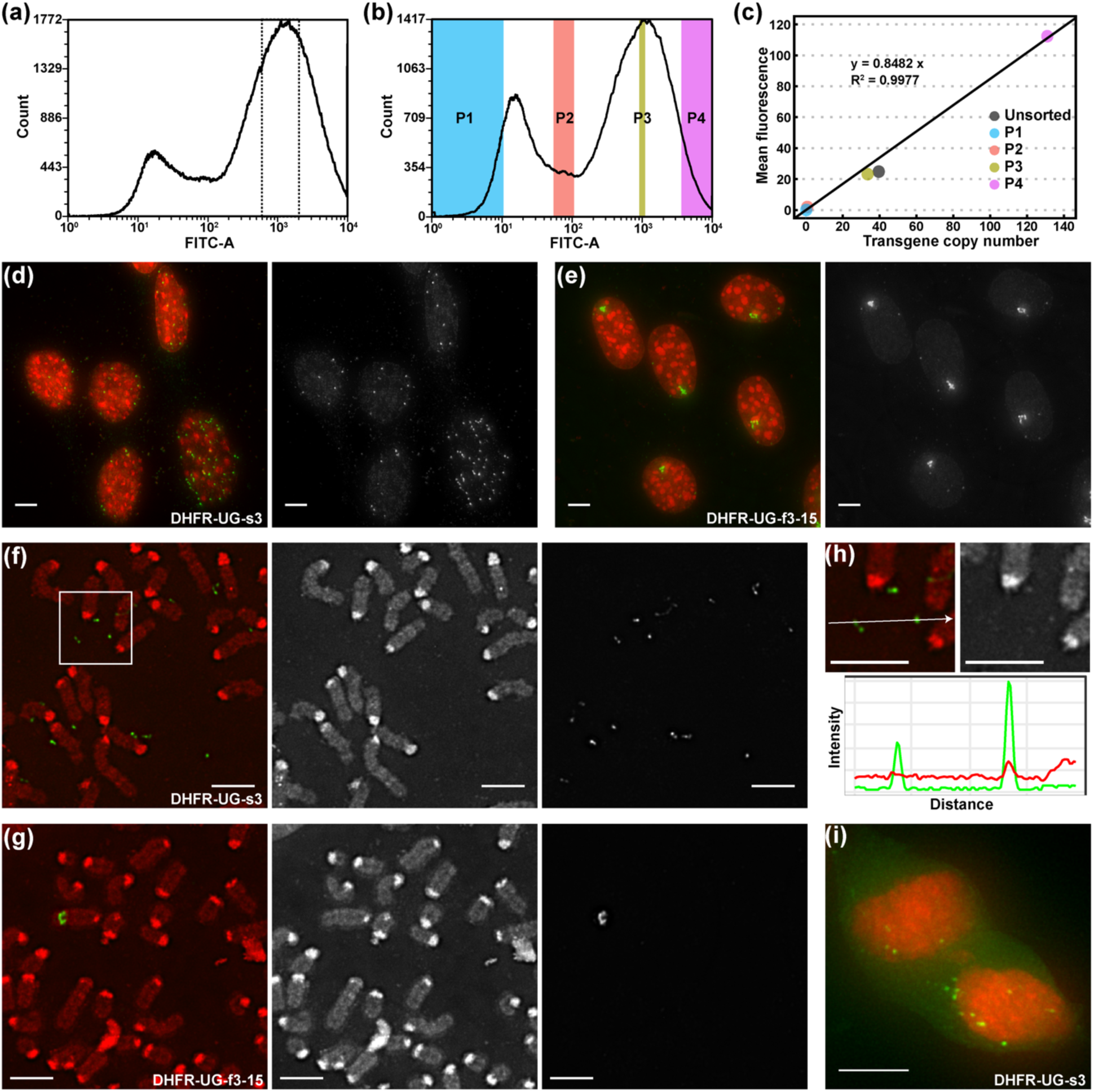
BAC transgenes exist as episomes in heterogeneously expressing clones. (a-c) BAC copy number analysis of sub-populations of a heterogeneous clone, DHFR-UG-s3, with different fluorescence levels. (a) GFP fluorescence histogram of DHFR-UG-s3 cells during first sorting (y-axis-cell number; x-axis-GFP fluorescence level). Cells within a narrow peak-window (dotted lines) were sorted by FACS. (b) GFP fluorescence histogram of sorted DHFR-UG-s3 cells after one week of cell growth. Cells within the four colored windows (P1-4) were sorted by FACS and used for BAC copy number estimation by qPCR. (c) Mean GFP fluorescence (y-axis) vs copy number (x-axis) of the four cell sub-populations and the original unsorted population shows linear correlation between fluorescence levels and copy number (R^2^=0.99). (d-e) DNA FISH over interphase nuclei of the heterogeneous clone DHFR-UG-s3 (d) and a uniform clone DHFR-UG-f3-15 (e) to visualize the BAC transgenes. Maximum-intensity projections are shown. Gamma=0.5 was applied to FISH channel after projection to better display low intensity FISH spots. (f-g) DNA FISH over mitotic spreads of the heterogeneous clone DHFR-UG-s3 (f) and the uniform clone DHFR-UG-f3-15 (g). (h) DAPI intensity over an episome with strong FISH signal and one with weak FISH signal. Top: enlarged view of the white square area in (f); bottom: DAPI (red) and FISH signal (green) intensity profile along the white arrow in the top panel. (i) A pair of telophase nuclei of the heterogeneous clone, DHFR-UG-s3, showing unequal segregation of episomal BAC transgenes during mitosis. (d-i) Red-DNA DAPI stain; green-BAC FISH signal. Scale bars = 5 μm.

To identify the source of this BAC copy-number instability, we next used DNA FISH to visualize BAC transgenes within interphase nuclei and mitotic chromosome spreads. We compared the distribution of BAC transgenes within the heterogeneous clone, DHFR-UG-s3, versus a uniform clone, DHFR-UG-f3-15.

DNA FISH suggested that whereas the uniform clone contained cells with an integrated DHFR BAC array, the heterogeneous clone contained cells in which the DHFR BAC was present as episomes. Specifically, interphase FISH against the DHFR BAC in the heterogeneous clone revealed multiple, noncontiguous, small spots distributed randomly throughout the nuclei (Figure 5d). The number of these spots was highly variable in different cell nuclei, suggesting unequal segregation of BAC transgenes. In contrast, most cells from the uniform clone showed just one large, fiber-like FISH spot per nucleus (Figure 5e). Moreover, FISH spots in mitotic spreads from the heterogeneous clone were either touching or spatially separated from the chromosomes (Figure 5f), whereas FISH spots in mitotic spreads from the uniform clone were always located within the chromosome (Figure 5g). The number of FISH spots per mitotic spread was highly variable in the heterogeneous clone, with each spot much smaller than the single FISH spot visualized within the mitotic chromosome from the uniform clone. Interestingly, the FISH spots in the heterogeneous clone mitotic spreads had weak DAPI staining, varying from slightly elevated over background to no difference from background (Figure 5h), suggesting these structures are much smaller than previously described double minute chromosomes (DMs) generated by gene amplification (96–98).

Using DNA FISH of both interphase nuclei and mitotic spreads, we confirmed this finding of integrated BACs in all uniformly expressing clones versus episomal BACs in all heterogeneously expressing clones in additional cell clones carrying BAC transgenes based on three different BAC scaffolds (Supplementary Figure S2 and data not shown). Specifically, this includes 4 heterogeneous and 4 uniform DHFR-UG BAC clones, 3 heterogeneous and 6 uniform HBB-UG BAC clones, and one heterogenous and 2 uniform COL1A1-UG BAC clones.

Unequal segregation of these BAC episomes during cell division would explain the heterogeneity of BAC transgene copy number in the cell population of heterogeneous clones, leading to variability of reporter gene expression. Indeed, telophase cells from heterogeneous clones showed unequal numbers of FISH spots in the two daughter nuclei (Figure 5i). In the absence of continued drug selection, we would expect cells that have lost BAC transgenes will accumulate if there is any selective growth advantage for cells with fewer BAC copies.

### BAC episomes are circular and ∼1 Mb in size

We analyzed the average amount of DNA per BAC episome, using two independent methods-light microscopy and pulsed-field gel electrophoresis (PFGE). Both methods produced a similar estimate of ∼800-1000 kb per BAC episome.

Using light microscopy, we measured the average DHFR BAC episome DAPI integrated staining intensity in mitotic spreads from cell clone DHFR-UG-s3 relative to the smallest mouse chromosome (chr19) with known DNA content of 61.4 Mbp (Figure 6a). This comparison produced an estimated mean episome size of 770 kb in this DHFR-UG-s3 clone.

**Figure 6.**
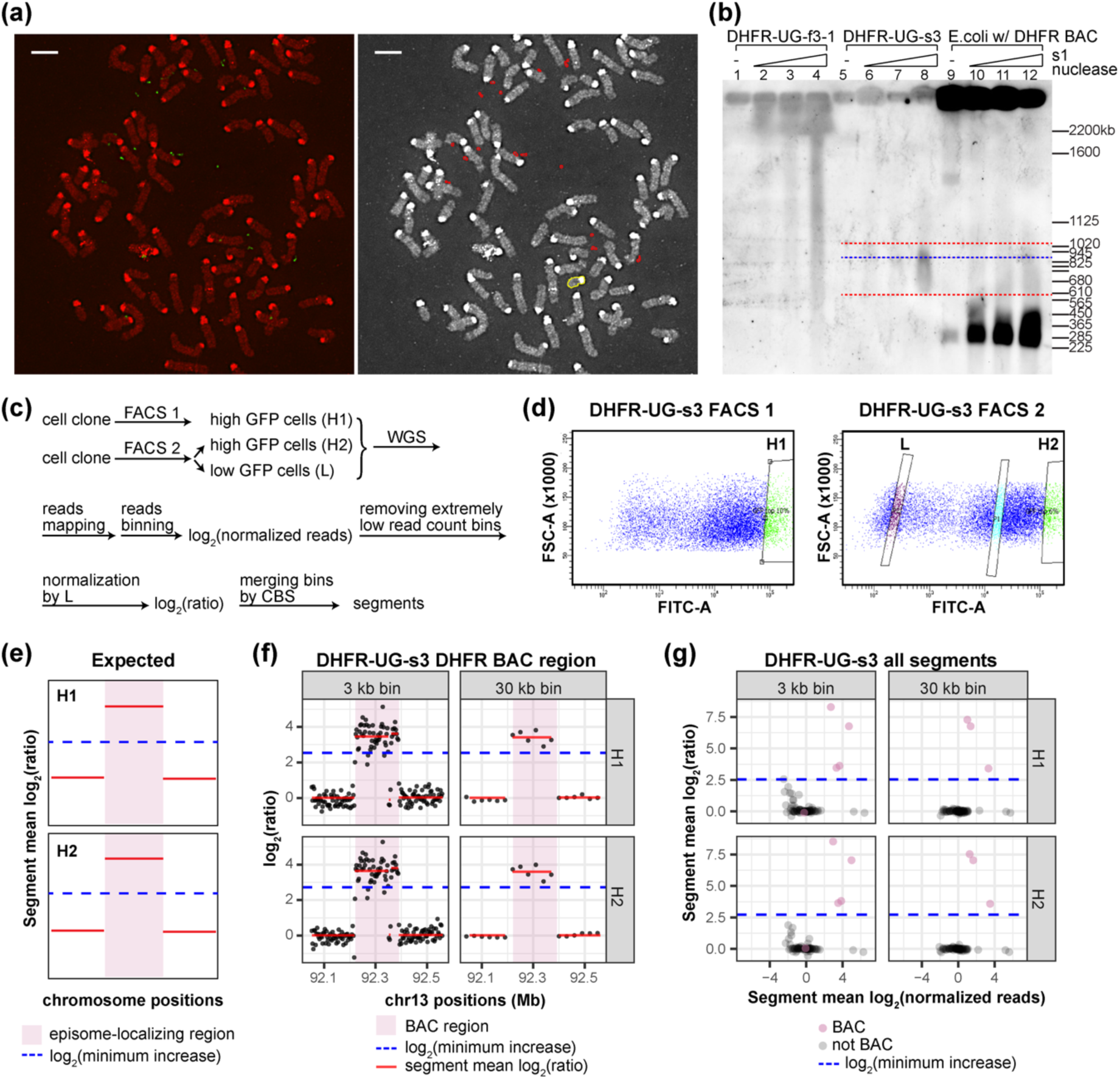
BAC episome size estimation and CNV analysis. (a) Estimation of average episome size in the DHFR-UG-s3 clone using mitotic FISH. Red-DNA DAPI stain; Green-BAC FISH signal; Red circles: regions of interest (ROIs) of FISH spots used for analysis; Yellow circles: ROI of the smallest chromosome in the field. Scale bars = 5 μm. This panel reuses the image in Figure 5f for analysis. (b) Southern hybridization using probes prepared from the DHFR BAC of cellular DNA without enzyme digestion, or digested with increasing amount of S1 Nuclease, separated by PFGE. Lane 1-4: uniform clone DHFR-UG-f3-1; Lane 5-8: heterogeneous clone DHFR-UG-s3; Lane 9-12: *E. coli* carrying the DHFR BAC. (c-g) CNV analysis of the DHFR-UG-s3 clone. (c) Flow chart of the CNV analysis. (d) Two FACS experiments for collecting cells with high (H1 and H2), and low (L) fluorescence subpopulation. x-axis-FITC channel intensity; y-axis-forward scatter; H1, H2, and L-sorting windows. (e) Episome-localizing genomic regions (pink highlighted regions) are expected to have mean log_2_(ratio) (red line) equal to or greater than log_2_(estimated minimum copy number increase) (blue dashed line). (f) log_2_(ratio) of individual bins (dark gray dots) and the segment mean log_2_(ratio) (red lines) around the *Dhfr-Msh3* locus belonging to the DHFR BAC (pink highlight) in the H1 and H2 subpopulations of the DHFR-UG-s3 clone. (g) Scatter plot of segment mean log_2_(ratio) vs segment mean log_2_(normalized reads) of all segments of the H1 and H2 subpopulations of the DHFR-UG-s3 clone. Pink dots-segments belonging to the DHFR BAC, including the *Dhfr-Msh3* locus, UBC-GFP-ZeoR and the BAC vector; Black dots- remaining segments in the genome. (f-g) Blue dashed line: log_2_(estimated minimum copy number increase).

Using PFGE, we observed that the BAC episomes were circular and estimated the modal BAC episome size to be ∼900 kb and 1 Mbp for DHFR-UG and HBB-UG BAC episomes in cell clones DHFR-UG-s3 and HBB-UG-100d3, respectively. Two different cell clones, DHFR-UG-f3-1 and HBB-UG-fD2, were used as negative controls as they contained the same DHFR-UG or HBB-UG BAC DNA as the cell clones with episomes but the BAC DNA was integrated within endogenous mouse chromosomes. *E. coli* strains containing the DHFR or HBB BACs were used as positive controls for detection of circular episomes.

Pulsed-field gels were analyzed by Southern blotting using pooled BAC DNA PCR products as the hybridization probes. Linear but not circular DNAs migrate in pulsed-field gels. Similar to the *E. col*i controls containing circular BACs, the Southern blot signals for the BAC DNA from the two clones containing episomes did not migrate out of the wells (Figure 6b and Supplementary Figure S3a-b), consistent with circular rather than linear BAC episomes.

To validate that the BAC episomes are really circular, and to estimate their size, the agarose-embedded DNA was digested using the ssDNA specific Nuclease S1 prior to PFGE and Southern blot hybridization. After removal of proteins, circular DNA episomes in both bacteria and mammalian cells are typically negatively supercoiled. This supercoiling generates torsional stress which is relieved by local formation of single-stranded regions. Thus, S1 nuclease has been used to cut these single-stranded regions and linearize circular DNA episomes (99–101). After S1 digestion, DNA from the cell clones carrying BAC episomes now showed DNA smears with peak intensities of ∼900 kb and 1 Mb for the DHFR-UG-s3 and HBB-UG-100d3 cell lines, respectively (Figure 6b and Supplementary Figure S3a-b). In contrast, after S1 nuclease digestion, DNA from the integrated BAC clones showed signals within the wells and above 2 Mb, overlapping the smears of fragmented genomic DNA (Figure 6b and Supplementary Figure S3a-b). DNA of *E. coli* containing DHFR BAC and HBB BAC episomes produced bands at ∼200-300 kb, in addition to signals in the wells (Figure 6b and Supplementary Figure S3a-b) after S1 nuclease digestion. These estimated BAC sizes measured slightly larger than the actual BAC sizes (∼200 kb), indicating there might be a slight overestimation of episome sizes using our PFGE running conditions.

### BAC episomes contain no detectable host DNA as revealed by CNV analysis

The propagation of BAC transgenes as episomes was unexpected. A major question is whether these episomes consist solely of BAC DNA, or whether host DNA is also included and possibly required for episome propagation.

We first compared the estimated episome DNA content size with estimates of BAC copies per episome. BAC episome sizes estimated by either light microscopy or PFGE were approximately twice as large as predicted from qPCR BAC copy number estimates (Table 2). The estimated average BAC content per episome was 445 kb in DHFR-UG-s3 and 716 kb in HBB-UG-100d3.

**Table 2.** BAC copy number, episome copy number, and BAC DNA content per episome in clone DHFR-UG-s3 and clone HBB-UG-100d3.

| Sample Name | BAC copy number per cell | Episome copy number per cell | BAC copy number per episome | BAC size (kb) | BAC content per episome (kb) |
| --- | --- | --- | --- | --- | --- |
| DHFR-UG-s3 | 15.4 | 6.2 (n=99) | 2.5 | 178 | 445 |
| HBB-UG-100d3 | 14.9 | 4.5 (n=100) | 3.3 | 217 | 716 |

The difference in the estimated BAC DNA content per episome and the average episome size is at most a few hundred kb, and may be accounted for by inaccuracies of the qPCR copy number, PFGE size estimation, and possible variation in sequence representation within the BAC episomes due to shearing of DNA and/or recombination during the transfection and creation of the BAC episomes.

Alternatively, this difference in episome size versus qPCR estimation of BAC copies per episome could also be caused by presence of host cell genomic DNA on the episomes. To search with higher sensitivity for the possible presence of host DNA within the episome, we performed Whole Genome Sequencing (WGS) based copy number variation (CNV) analysis of the two clones, DHFR-UG-s3 and HBB-UG-100d3. Genomic regions present on the episomes would appear amplified in cells containing episomes (test sample), comparing to cells with no episomes (reference sample). Thus the ratio in the number of reads for a given bin between the test sample and the reference sample was calculated. To reduce noise, bins were merged into segments based on the log_2_(ratio), using a circular binary segmentation (CBS) algorithm (78, 79). The mean log_2_(ratio) of each segment was used to estimate the CNV of this segment in the test sample relative to the reference sample.

Mouse 3T3 cells show genomic instability; therefore we anticipated CNV between the parental cell line and individual clones. To reduce false-positives derived from CNV between different 3T3 clones, independent of episome content, we used cells with low reporter gene fluorescence sorted from the cell clone containing the episomal BAC transgenes (region L, Figure 6c-d, and Supplementary Figure S4a) as the reference sample. To further reduce false positives, we also imposed constraints that copy number increase for true positive regions should be reproducible between experimental replicates and correlate with episomal copy number. We calculated the estimated CNV in cells sorted with high (H2) reporter gene expression, using sorted cells with low (L) expression as the reference sample (H2, L, Figure 6c-d, and Supplementary Figure S4a). We also compared the estimated CNV in cells sorted with high reporter gene expression in an independent experiment (H1, Figure 6c-d, and Supplementary Figure S4a) with the estimated CNV from the first experiment. All samples were sequenced to ∼2x coverage.

We used 3 and 30 kb bin sizes for analysis. To reduce noise, we excluded all bins in the test sample with zero reads (5.5% of total bins for 3 kb bin analysis and 3.5% for 30 kb bin analysis) plus extreme outlier bins, defined by the lower quantile minus 4 times the interquantile distance, with unusually low read count (∼0.5% of total bins for 3 kb bin analysis and 2.5% for 30 kb bin analysis) in the test sample before calculating ratios.

As a test of our analysis method, we compared the mean segment log_2_(ratio) of the BAC regions in H1 and H2, generated by the above analysis method, to the fold increase of BAC regions in H1 and H2 relative to L measured by qPCR. As expected, the results from the CNV and qPCR analysis were very similar (Supplementary Figure S5).

We were interested in asking whether a specific host DNA element was present on each episome copy present within a cell clone. We estimated that on average the sorted cells with high reporter gene expression had 15-20 episome copies per cell, depending on the cell clone, based on qPCR of BAC DNA sequences and the estimated number of BACs per episome (Table 3).

**Table 3.** BAC copy number, estimated episome copy number, and estimated minimum copy number increase of episome-localizing DNA in H1 and H2 subpopulations relative to L subpopulation of clone DHFR-UG-s3 and clone HBB-UG-100d3.

| Sample Name | BAC copy number per cell | Estimated episome copy number per cell | Minimum copy number increase of episome-localizing DNA relative to L |
| --- | --- | --- | --- |
| DHFR-UG-s3_H1 | 49.4 | 19.8 | 5.8 |
| DHFR-UG-s3_H2 | 57.5 | 23.1 | 6.6 |
| DHFR-UG-s3_L | 0.3 | 0.1 | / |
| HBB-UG-100d3_H1 | 48.5 | 14.7 | 4.6 |
| HBB-UG-100d3_H2 | 51.2 | 15.5 | 4.8 |
| HBB-UG-100d3_L | 0.2 | 0.1 | / |

We estimated theoretical minimum copy number increase for episome-localizing host DNA (minimum increase) in the H1 and H2, based on BAC copy number measured by qPCR, and assuming the NIH 3T3 to be tetraploid and each episome to have the same host cell genomic DNA (Table 3). Segments with mean log_2_(ratio) equal to or greater than log_2_(minimum increase) in both H1 and H2 samples were selected as candidates for being on the episomes (Figure 6e).

Interestingly, all candidate segments identified belonged to the BAC regions, including the UBC-GFP-ZeoR and the BAC vector (Figure 6f-g, Supplementary Figure S4b-c and Supplementary Figure S6), and no other mouse genomic sequence satisfied all of the above conditions.

In conclusion, we could not detect host cell DNA reproducibly present on all episomal copies using bin sizes of either 3 or 30 kb. We therefore conclude BAC DNA itself is sufficient for the creation and propagation of these BAC episomes. We cannot exclude the possibilities, however, that an unmappable, repetitive host DNA sequence is present on the episomes and confers their ability to propagate or that different host DNA sequences are present on each episome present within a single cell clone.

### Multiple promoters added to BACs support formation of episomal BAC transgenes but only in certain cell lines

Because we did not observe episomal BAC transgenes in our original BAC-TG EMBED work using the CMV-mRFP-SV40-ZeoR reporter gene (51), we hypothesized that addition of the UBC-GFP-ZeoR reporter gene might be responsible for BAC episome formation. Our dual-reporter assay showed that the UBC promoter was much stronger than the CMV promoter; therefore, we further hypothesized that promoter strength might correlate with the frequency of BAC episome formation.

To test this hypothesis, we isolated clones stably transfected with the dual-reporter DHFR BAC transgenes and examined reporter gene expression patterns in these clones by flow cytometry (Supplementary Figure S7a). As expected, no heterogeneously GFP/RFP expressing clones where observed when the mRFP reporter gene was driven by CMV promoter (n=13) or B2M promoter (n=6). In contrast, we observed ∼70% or ∼30% heterogeneously GFP/RFP expressing clones when the mRFP was driven by the EEF1a promoter (12/18) or the RPL32 promoter (10/29), respectively (Supplementary Figure S7a-b). We confirmed that BAC transgenes in these heterogeneously expressing clones were episomal using DNA FISH (Supplementary Figure S7c).

These results using human promoter sequences added to the BACs, did show a rough correlation of promoter strength with the frequency of clones containing episomes. However, when we examined a series of DHFR BAC constructs, we instead observed clones with episomes using BAC transgenes containing the dual reporter, selectable marker CMV-mRFP-SV40-ZeoR reporter cassette. This included the identical DHFR BAC construct used in our previous BAC-TG EMBED work (51), as well as various DHFR BAC deletions (Supplementary Figure S8a). All the DHFR BAC constructs contain LacO repeats, and a NIH 3T3 derived clone expressing EGFP-LacI was used for transfection, so that BAC transgenes could be observed directly in fixed cells. Although all clones showed a unimodal flow cytometry expression pattern, explaining why we did not observe this phenomenon previously, a large fraction (DHFR-c27: 1/2, DHFR-c27d2: 2/10, DHFR-c27d3crz: 2/16, DHFR-c27d4: 7/10) of clones showed episomal BAC transgenes (Supplementary Figure S8b-c).

Thus, the promoter used to drive reporter and/or selectable markers appears to determine not whether episomal BAC transgenes are established but rather whether a unimodal versus bimodal distribution is observed in cells containing these episomal BAC transgenes. The presence of strong promoters (UBC, EEF1a and RPL32) appears to allow the formation of bimodal distributions of reporter gene expression, possibly related to the balance between the degradation rate of the initially high levels of selectable marker versus the rate of loss of BAC transgene episomes during cell division.

Next, we tested whether BACs can form episomes in a different cell line other than mouse NIH 3T3 fibroblasts. Previously, we observed cell clones containing only integrated BAC transgenes in CHO (93, 95) and mouse ES cells (54, 94). Reasoning that cancer cells with some level of genomic instability might be more prone to formation of BAC episomes, we tested the human colorectal carcinoma epithelial cell line, HCT116, using the 2207K13-UG BAC which produced 79% episome clones in NIH 3T3 cells.

Four out of 32 stable clones showed a heterogeneous GFP distribution similar to that observed in NIH 3T3 episome clones, with a broad high fluorescent peak and a tail/secondary peak near the auto-fluorescence level (Supplementary Figure S9). However, none of these four clones showed episomal BAC transgenes by DNA FISH (Supplementary Figure S10a). Instead, most cells in each clone showed the same number (one or two) of spots, but these spots varied in size from cell to cell. Therefore, it appears that the broad GFP peaks in these four clones are due to some form of genomic instability leading to CNV of integrated transgene arrays. Interestingly, one clone, HCT116-k13_06, out of the 32, which had a single GFP peak, showed a small fraction of cells of with episomal BAC transgenes, in contrast to the vast majority of cells which contained integrated BAC transgenes (Supplementary Figure S10b). One out of 24 subclones of this HCT116-k13_06 clone, HCT116-k13_06-10, showed a similar mixed population with either integrated BACs or episomal BACs, similar to the parent clone HCT116-k13_06 (Supplementary Figure S10c). The low frequency of clones with episomal BACs, the variable size of the integrated BAC transgene arrays, and the co-existence of integrated and episomal BAC transgenes in the same cells and from the same clone suggests these episomes might arise from the well-known phenomenon of gene amplification (96–98).

Similarly, a small percentage of clones carrying the GAPDH BAC in stable mouse ES cell colonies showed broad GFP expression peaks by flow cytometry, but FISH revealed this was due to variable size, integrated BAC transgene arrays (Binhui Zhao, Ph.D thesis), due presumably to some type of CNV induced by genomic instability of these transgene arrays.

In conclusion, the high frequency establishment of BAC transgene episomes seen in mouse 3T3 cells does not appear to occur in either HCT116 or mouse ES cells, or at detectable frequency in CHO cells (54, 93–95).

### Expression of multiple-reporters by BAC-MAGIC

As a proof-of-principle application of our improved toolkit for BAC TG-EMBED, we created a multi-transgene BAC to label simultaneously the nuclear lamina, nucleoli, and nuclear speckles with a single stable transfection. The original DHFR BAC was used for this multi-transgene expression. A SNAP-tagged Lamin B1 reporter mini-gene was used to label the nuclear lamina, a SNAP-tagged Fibrillarin the nucleoli, and an mCherry-Magoh the nuclear speckles. We used the RPL32 promoter to drive the expression of the SNAP-tagged Lamin B1, and a promoter of intermediate strength, PPIA, for the SNAP-tagged Fibrillarin and the mCherry-tagged Magoh, which are both abundant proteins.

Previously, we used random Tn5 transposition to introduce expression cassettes into BAC scaffolds (50), but this approach is limited in the number of serial insertions that can be made due to the remobilization of existing transposons, its requirement for multiple selectable markers, and the randomness of the insertion sites. Alternatively, BAC recombineering using antibiotic resistance genes as positive selectable markers have been used to insert expression cassettes into precise locations on the BACs. However, like transposition, this method relies on the availability of multiple selectable markers and introduces unwanted selectable markers into the BACs. An alternative BAC recombineering scheme using cycles of *galK*-based positive selection to insert sequences followed by negative selection to remove *galK* have been used to make multiple BAC modifications without addition of unwanted selectable markers. However, the low efficiency of negative selection, due to a high background of competing, spontaneous deletions of mammalian DNA with its high repetitive DNA content, makes this approach quite time and labor intensive. Typically, one month is required for each cycle of insertion of DNA by positive selection, removal of the selectable marker by negative selection, and subsequent screening and testing of DNA from colonies that survive the negative selection to identify the small fraction of colonies containing the desired homology-driven, specific deletion of just the selectable marker.

To accelerate creation of BACs containing multiple transgene, we created a new BAC assembly approach, BAC MAGIC (**<u>BAC</u>**-**<u>M</u>**odular **<u>A</u>**ssembly of **<u>G</u>**enomic loci **I**nterspersed **<u>C</u>**assettes). BAC MAGIC combines the DNA assembler method in yeast (81, 82) and/or Gibson assembly (80) with traditional cloning methods to create a number of BAC recombination modules followed by sequential rounds of BAC recombineering in which one fragment is inserted using one selectable marker followed by addition of a new fragment overlapping the previous fragment using a second positive selectable marker which replaces the first (102). Each round of fragment insertion only requires ∼ 1 week for transformation and screening of clones. In this way, 45 kb of the DHFR BAC was effectively reconstructed such that DHFR sequences remained but 3 fluorescent mini-gene expression cassettes were added, each spaced by ∼10 kb of DHFR sequence (Figure 7a-b, Supplementary Figure S11). The large homologous sequences flanking each expression cassette reduces recombination between similar sequences in other expression cassettes already inserted into the BAC, increasing the efficiency of this overall approach.

**Figure 7.**
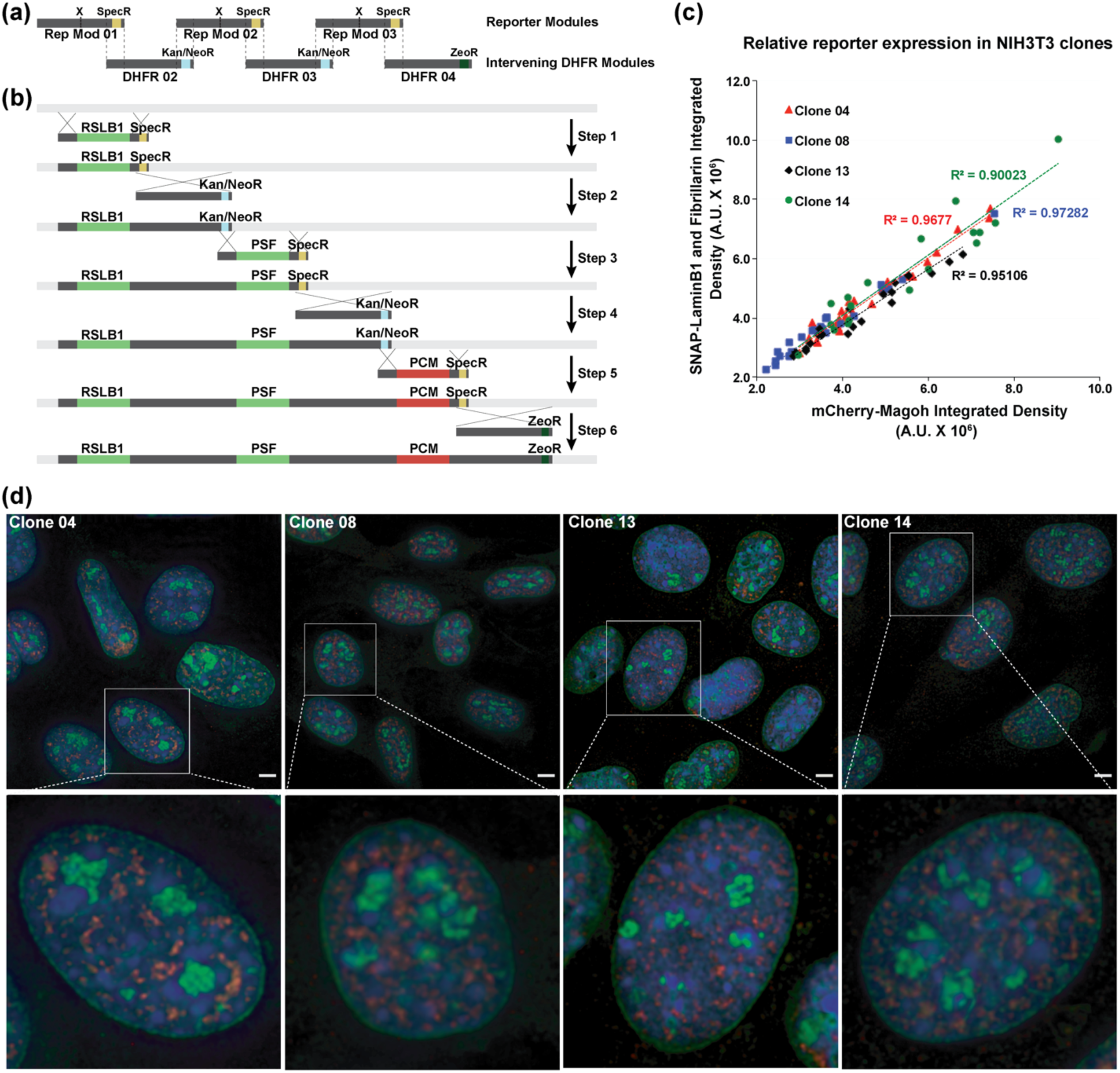
BAC-MAGIC and simultaneous multi-reporter expression. (a-b) Construction of the multi-reporter DHFR BAC by BAC-MAGIC. (a) Modular design of BAC-MAGIC. Reporter module 01, 02 and 03 contain reporter gene expression cassettes (X), DHFR BAC homologous sequences (dark gray), and Spectinomycin resistance markers (SpecR, yellow) near the 3’ ends for bacterial selection. Intervening DHFR module 02, 03 and 04 contain DHFR BAC homologous sequences (dark gray), and antibiotic resistance markers near the 3’ ends (Kanamycin/Neomycin resistance marker (Kan/NeoR, blue) in module 02 and 03 for bacterial selection, and Zeocin resistance marker (ZeoR, dark green) in module 04 for dual selection in bacterial or mammalian cells). The dotted lines mark homologous regions between the reporter modules and the intervening DHFR modules. (b) Six sequential steps of BAC recombineering introduce three reporter expression cassettes, RPL32-driven SNAP-tagged Lamin B1 (RSLB1), PPIA-driven SNAP-tagged Fibrillarin (PSF), and PPIA-driven mCherry-Magoh, onto the DHFR BAC (light gray) with ∼10 kb of intervening DHFR BAC sequences (dark gray). Homologous regions are indicated by crossed lines. (c) Relative expression of the SNAP-tagged Lamin B1 and Fibrillarin to the mCherry-Magoh reporter in four representative NIH 3T3 cell clones (04, 08, 13 and 14) containing the multi-reporter BAC. Integrated fluorescence intensities per cell of SNAP--fluorescein (y-axis) and mCherry-Magoh (x-axis) are plotted. Linear regression lines (y-intercepts set to 0) are shown with corresponding R-squared values. Number of nuclei of each clone analyzed range from 18 to 27. Red-Clone 04; Blue-Clone 08, Black-Clone 13; Green-Clone 14. (d) Representative images (maximum intensity projections of 2-3 optical sections) from the four cell clones (Clone 04, 08, 13 and 14) showing expression of the three reporter genes. Nuclear lamina is labeled with SNAP-tagged Lamin B1 (green), nucleoli with SNAP-tagged Fibrillarin (green), and speckles with mCherry-Magoh (red). One magnified nucleus from each representative field (top panel) is shown in the bottom panel. Scale bars = 5 μm.

We began the process using a DHFR BAC. After six rounds of BAC recombineering, we had created a BAC with four expression cassettes (Figure 7b): a SNAP-tagged Lamin B1 minigene, a SNAP-tagged Fibrillarin minigene, a mCherry-Magoh minigene, and a ZeoR selectable marker.

We tested simultaneous expression of the three reporters in 17 independent NIH 3T3 cell clones transfected with the multi-reporter BAC by examining fluorescence in fixed cells under a microscope (SNAP-tagged proteins were labeled with a Fluorescein conjugated SNAP tag substrate before fixation). We observed uniform expression of all the three reporters in 16/17 clones. The loss of SNAP-Lamin B1 expression in one of the clone (Cl#16) may be due to random breakage of the BAC during transfection, as PCR revealed the absence of this minigene from the cell clone. Similarly, 12/14 U2OS human osteosarcoma cell clones showed both SNAP-Lamin B1 and SNAP-Fibrillarin expression after transfection of a BAC containing only these two expression cassettes (data not shown).

Within individual cells, a linear correlation was observed between the integrated fluorescence intensity per cell of SNAP-tagged proteins Lamin B1 and Fibrillarin versus mCherry-Magoh in 4/4 representative NIH 3T3 clones (04, 08, 13 and 14, Figure 7c). Moreover, these fluorescently tagged proteins showed uniform rather than variegating expression in different cell nuclei of the same clone observed under the microscope (Figure 7d).

## DISCUSSION

We previously demonstrated the utility of the BAC TG-EMBED method to achieve position-independent, copy-number-dependent, one-step transgene expression in mammalian cells (51, 54). Here, we have extended the BAC TG-EMBED methodology through four new advances and provided a proof-of-principle demonstration of this new methodology by efficiently creating cell lines stably expressing uniform levels of three different fluorescently tagged proteins-Lamin B1, Fibrillarin, and Magoh in a single stable transfection.

First, we describe a toolkit of endogenous promoters providing an ∼500-fold range in promoter strength varying from ∼5 fold higher to ∼100-fold weaker than the commonly used viral CMV promoter. As these promoters are from human genes shown to be expressed in a wide range of cell lines and tissues (60, 84–90), we expect them to support transgene expression in most cell types and independent of cell proliferation or differentiation state. While most of the previous studies on transgene promoters focused on conventional, strong promoters (53, 57–60), including a similar approach that expressed mini-genes within BAC scaffold (53), we included moderate-strength and weak promoters in our survey. The weak promoters we identified, such as GUSB and RPS3A, could possibly replace the commonly used minimal promoters or inducible promoters where a sustained low-level of transgene expression is needed. Moreover, this wide range of promoter strengths allows reproducible expression of multiple transgenes over a wide range of relative expression levels from a single BAC scaffold, lending itself to such purposes, for example, as the design of synthetic gene circuits, which typically requires expression of different components at reproducible relative expression levels (25).

Second, we show that with the UBC-GFP-ZeoR reporter gene, our BAC-TG EMBED system achieved stable reporter gene expression of integrated BAC transgenes for several months in the absence of drug selection. This is an improvement over the 30-80% drop in expression observed originally with the CMV-mRFP-SV40-ZeoR reporter gene (51). Both the UBC promoter and the CpG free GFP-ZeoR gene body could be contributing to this improvement.

Third, we show that at least with the UBC-GFP-ZeoR expression cassette, our BAC TG-EMBED system is not dependent on BAC scaffolds containing active DNA genomic regions but also works with BAC scaffolds containing silenced DNA genomic regions as well as gene deserts. UBC may represent a member of a class of active, house-keeping gene promoters that is relatively insensitive to chromosome position effects. This allows choice of a BAC scaffold for the BAC TG-EMBED method that will not co-express any genes other than the introduced transgene cassettes. In contrast, both our previous BAC TG-EMBED studies (51, 54) and similar work from other laboratories (52, 53), used only BACs containing highly-transcribed house-keeping genes, due to the assumption that either an active chromatin region or active 5’ *cis*-regulatory regions would be required for creating a transcriptionally permissive environment for transgene expression. Integration of the UBC-GFP-ZeoR reporter gene into the BAC was required for position-independent, copy-number dependent expression, as its expression was copy-number independent when the same UBC-GFP-ZeoR reporter gene was stably transfected by itself into cells. The expression levels of this UBC-GFP-ZeoR were similar, per copy number, in cell clones with episomal BAC transgenes to levels in clones with integrated BACs.

Fourth, we describe an episome version of our BAC-TG EMBED system. In a single experiment, clones containing either stably integrated or extrachromosomally maintained BAC transgenes can be isolated. Most of the widely used episomal vectors are either based on viral sequences or derived from the non-viral pEPI plasmid (38–41). A notable feature of the episomes generated by our BAC TG-EMBED system is that they are lost rapidly in the absence of drug selection, whereas both of the other two systems show selection-independent mechanisms for stable episome maintenance (43–47, 103). While episome stability is valuable for certain applications, in other cases one would like to be able to easily eliminate the episomes as needed. Moreover, in contrast to the low copy number of episomes per cell produced using the other two methods, the BAC TG-EMBED method yields tens of BAC copies per cell allowing for much higher transgene expression levels. Additionally, the sizes of the episomes generated by the BAC TG-EMBED method are much larger than those generated by the other two methods. In the two clones we examined, the episomes were ∼1 Mb and containing several copies of the BACs per episome and no detectable host DNA.

The high frequency creation and simple composition of these BAC episomes contrasts with human artificial chromosomes (HACs), which are special episomes, usually 1-10 Mb in size containing centromeric repeat sequences, mitotically stable, and maintained at low copy number (104–107). Capable of introducing large DNA sequences into recipient cells, HACs have shown great potential in a wide range of applications, such as recombinant protein production, drug selection and gene therapy (108–111). However, the construction of HACs remains non-trivial: it requires cloning of either telomere sequences and/or alphoid DNA, the formation of HACs occurs at very low frequency and only in certain cell lines (112), and the transfer of HACs from donor cells into recipient cells is difficult (113, 114). Moreover, the presence of large telomere sequences and/or alphoid DNA on the HACs, and the heterchromatic state associated with these repeats, increases the likelihood of transgene silencing.

In contrast, with our BAC-TG EMBED system, 10s-100s of stable cell clones containing multiple copies of ∼Mb-size episomes, likely containing only BAC DNA, can be obtained from a single transfection. Cells containing high copy numbers of these BAC episomes can be enriched by flow sorting, while cells from these clones containing no BAC episomes can be recovered after removal of drug selection and/or flow sorting. We anticipate that with additional engineering, these BAC episomes might possibly become a high-capacity episome system complementary to HACs, assuming they can be isolated from one cell line and then introduced and propagated in other cell lines.

It remains unclear how these BAC episomes form in NIH 3T3 cells and why they do not do so in other cell lines. In the two clones we studied, the episomes were circular DNA and composed of several BAC copies. Interestingly, previous studies have shown that plasmids containing a MAR that is also a replication initiation region (IR) could initiate gene amplification in certain primary cancer cells, forming homogenously staining regions (HSRs), integrating into existing double minutes (DMs) or forming DMs *de novo* in cells without DMs (115, 116). It is believed that the IR/MAR plasmids are initially replicated as extrachromosomal circles, and then they multimerize into larger circular molecules. These amplified circles further multimerize to form DMs, recombine with pre-existing DMs or integrate into chromosomes and initiate HSR formation (117, 118). This model is very similar to the episome model of gene amplification, where instead of the IR/MAR plasmids, small extrachromosomal circular DNAs, which are several hundred kb in size and are possibly produced by small chromosome deletions, initiates DM and HSR formation (97, 98, 119).

Given that both the MAR and IR sequences are ubiquitous in the mammalian genome, it is likely that the BACs used in this study also contain MAR and/or IR sequences. However, unlike the MAR/IR plasmids, these BACs did not generate typical HSRs when integrated into the chromosomes, and the episomes were much smaller than DMs in NIH 3T3 cells. One possible explanation is that the BACs undergo initial steps of gene amplification to form the episomes in NIH 3T3 cells, but the cells have mechanisms to stop the episomes from further multimerization or amplification. As gene amplification happens only in cancer cells, perhaps BACs can only form episomes in certain cell lines. As shown here, BAC transgene formed episomes in a small fraction of HCT116 cells, which could not be stably maintained even with drug selection. Further study of this BAC episome phenomenon may provide new insights into the process of gene amplification.

Alternatively, the formation of BAC transgene episomes in NIH 3T3 cells might occur through a process completely unrelated to gene amplification. Future work will be needed to determine the actual mechanism of this BAC episomal formation in mouse 3T3 cells.

To facilitate the assembly of BACs expressing multiple mini-genes, we developed BAC-MAGIC, allowing creation of a multi-transgene expressing BAC in several weeks, rather than the 4-5 months which would have been required by multiple rounds of DNA insertion using conventional BAC recombineering. Initial attempts to reassemble large, ∼50kb regions of DHFR using yeast DNA assembly failed, apparently due to recombination between repetitive elements within the DHFR BAC sequence as well as the expression cassettes. In contrast, assembly of 10-15 kb modules from several DNA fragments using yeast DNA assembly worked with high efficiency. BAC-MAGIC exploits Gibson and yeast DNA assembly to build smaller modules with efficient serial BAC recombineering to reconstruct large BAC constructs containing multiple mini-gene expression cassettes. More generally, BAC-MAGIC should provide a tool for reconstruction of large eukaryotic DNA sequences containing high numbers of repetitive elements.

Finally, as a demonstration of our new version of BAC TG-EMBED system, we created cell lines expressing three different fluorescently tagged proteins in a single stable transfection step requiring just several weeks to isolate and expand cell clones. Most cell clones expressed all three tagged proteins at uniform levels and at reproducible relative levels of expression. This contrasts with the 6-12 months we have devoted in previous studies to create similar cell lines expressing multiple tagged proteins (120) through a series of individual transfections followed by extensive screening of colonies to identify the small fraction expressing suitable levels of tagged proteins with minimal variegation and/or progressive long-term transgene silencing over time.

We anticipate that our expanded BAC TG-EMBED toolkit similarly will facilitate a wide range of applications requiring simultaneous expression of multiple transgenes.

## ACKNOWLEDGEMENTS

We thank Edith Heard (Curie Institute) for providing DHFR BAC (clone 057L22 from CITB mouse library), Veena K Parnaik (CSIR-CCMB, Hyderabad, India) for GFP-Lamin B1 plasmid, Miroslav Dundr (Rosalind Franklin University of Medicine and Science) for GFP-Fibrillarin plasmid, Huimin Zhao (University of Illinois Urbana-Champaign) for pQCXIN-TetR-mCherry plasmid, Peter Adams (Sanford Burnham Prebys Medical Discovery Institute) for BJ-hTERT cells, and KV Prasanth (University of Illinois Urbana-Champaign) for mRFP-Magoh plasmid. Use of the BD FACS AriaII was assisted by the flow cytometry facility stuff at Roy J. Carver Biotechnology Center, University of Illinois at Urbana-Champaign (UIUC).

