## Supplemental Data for "Versatile multi-transgene expression using improved BAC TG-EMBED toolkit, novel BAC episomes, and BAC-MAGIC"

Figure S1

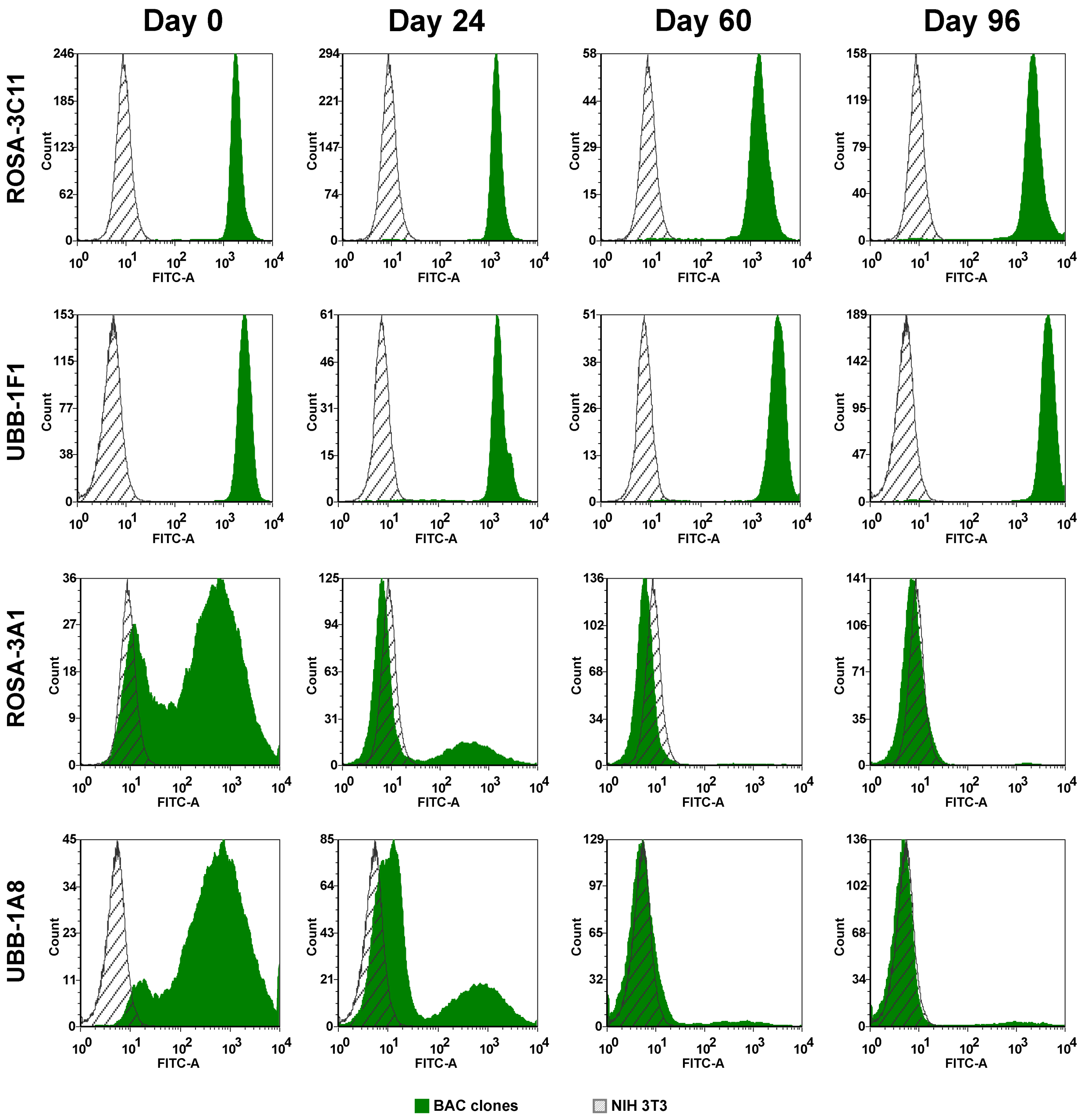
Day 0
Day 24
Day 60
Day 96
ROSA-3C11
UBB-1F1
ROSA-3A1
UBB-1A8
Count
FITC-A
BAC clones
NIH 3T3

Figure S2

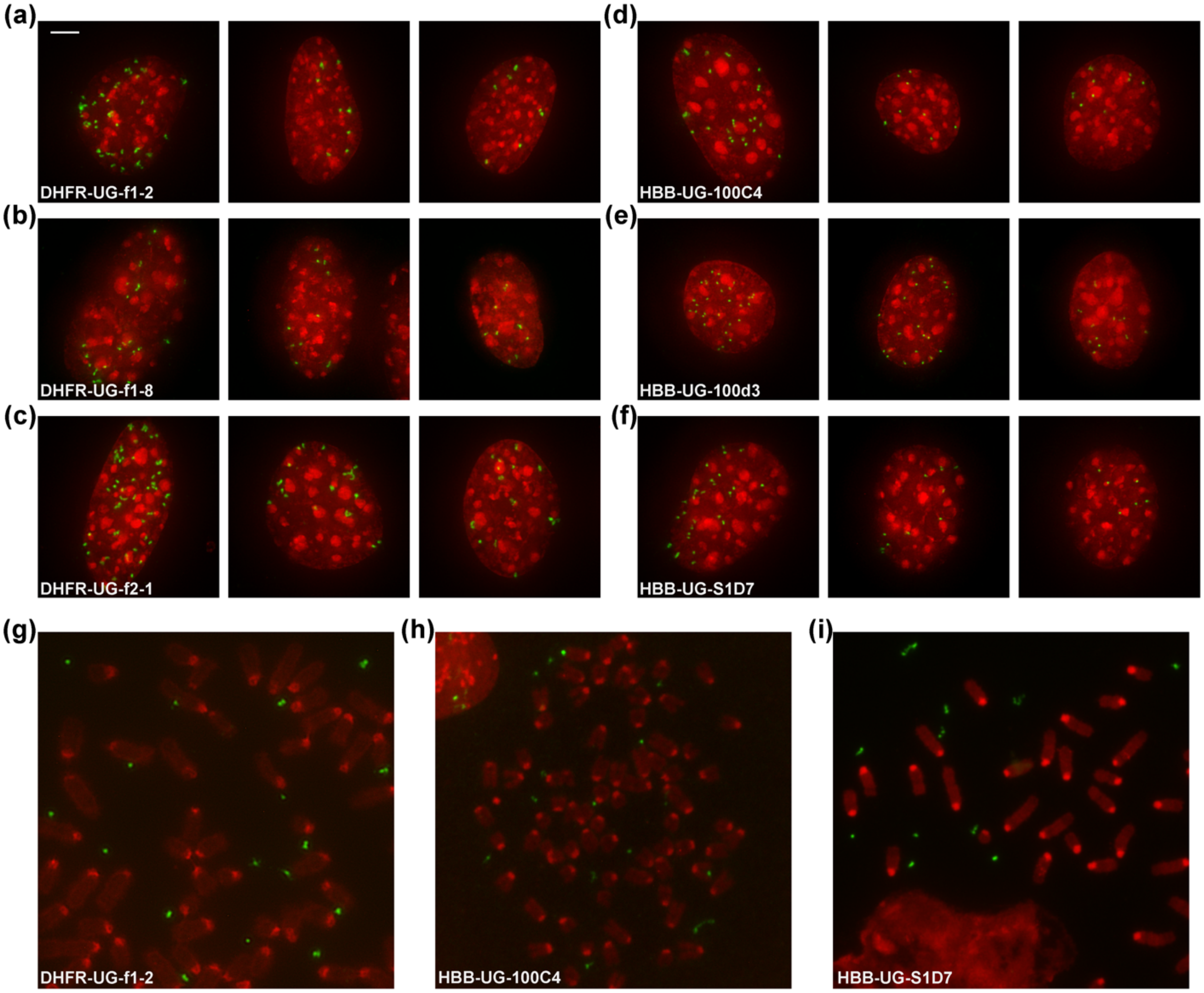
(a)
DHFR-UG-f1-2
(b)
DHFR-UG-f1-8
(c)
DHFR-UG-f2-1
(d)
HBB-UG-100C4
(e)
HBB-UG-100d3
(f)
HBB-UG-S1D7
(g)
DHFR-UG-f1-2
(h)
HBB-UG-100C4
(i)
HBB-UG-S1D7

Figure S3

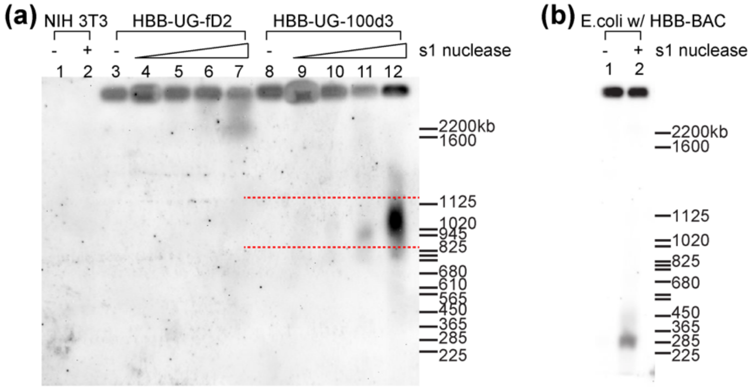


Figure S4

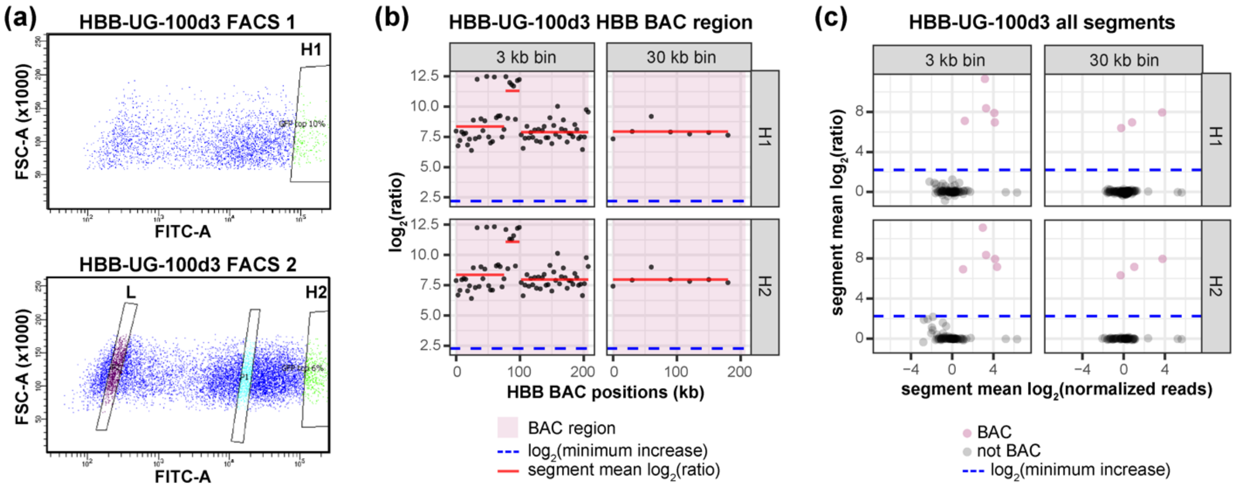

Figure S5

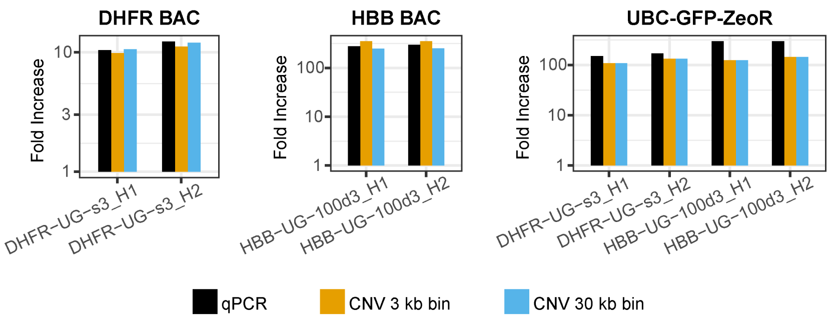


Figure S6

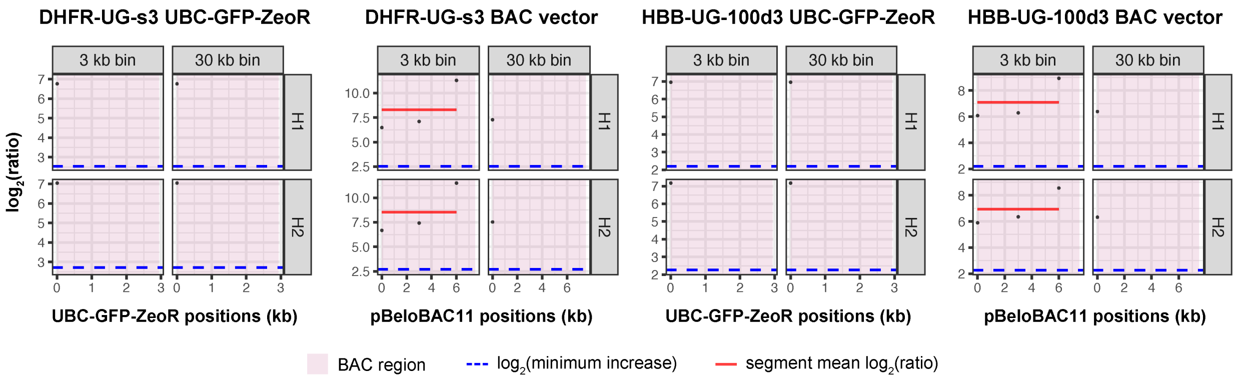

Figure S7

(a)

| Reporter Construct | Heterogeneous Clones% | Relative Promoter Strength |
|---|---|---|
| CMV–mRFP–SV40–ZeoR | 0 (n=13) | 1.0 |
| B2M–mRFP–SV40–ZeoR | 0 (n=6) | 0.3 |
| UBC–GFP–ZeoR | 60% (n=30) | 3.0 |
| UBC–mRFP–SV40–ZeoR | 45% (n=20) | 3.0 |
| RPL32–mRFP–SV40–ZeoR | 35% (n=29) | 5.5 |
| EEF1α–mRFP–SV40–ZeoR | 67% (n=18) | 4.6 |

(b)

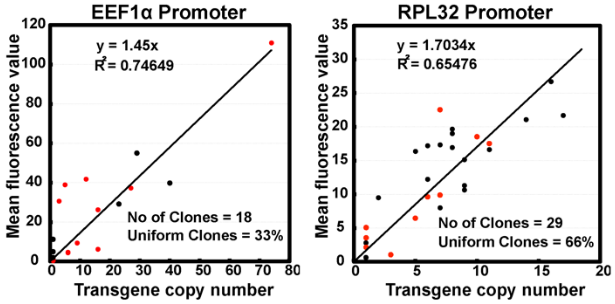


(c)

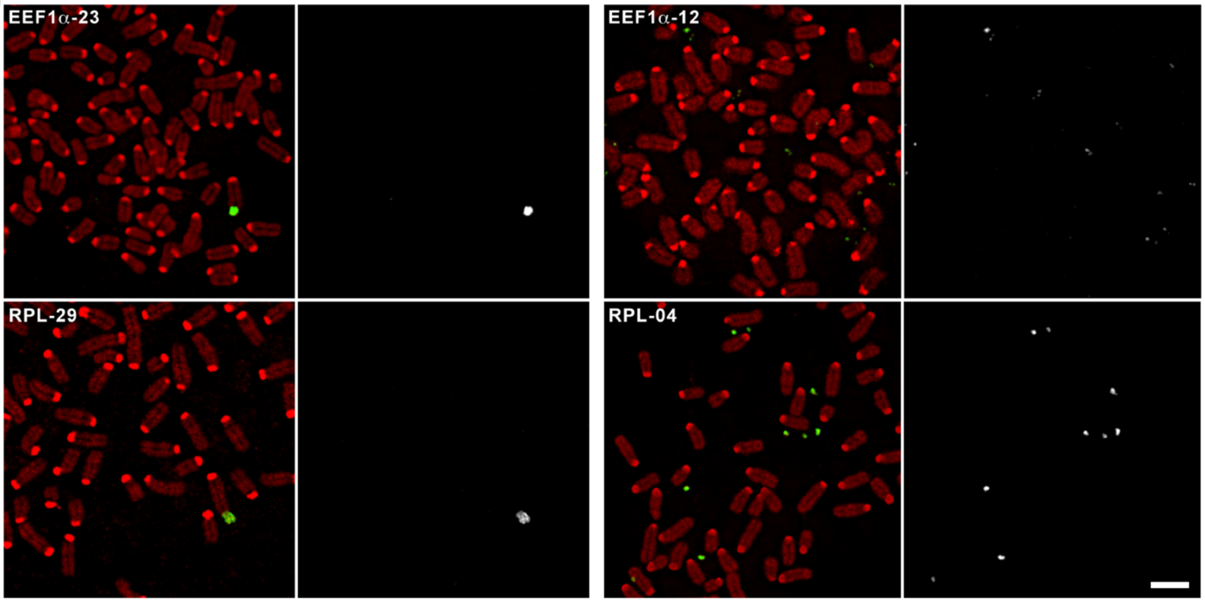

Figure S8

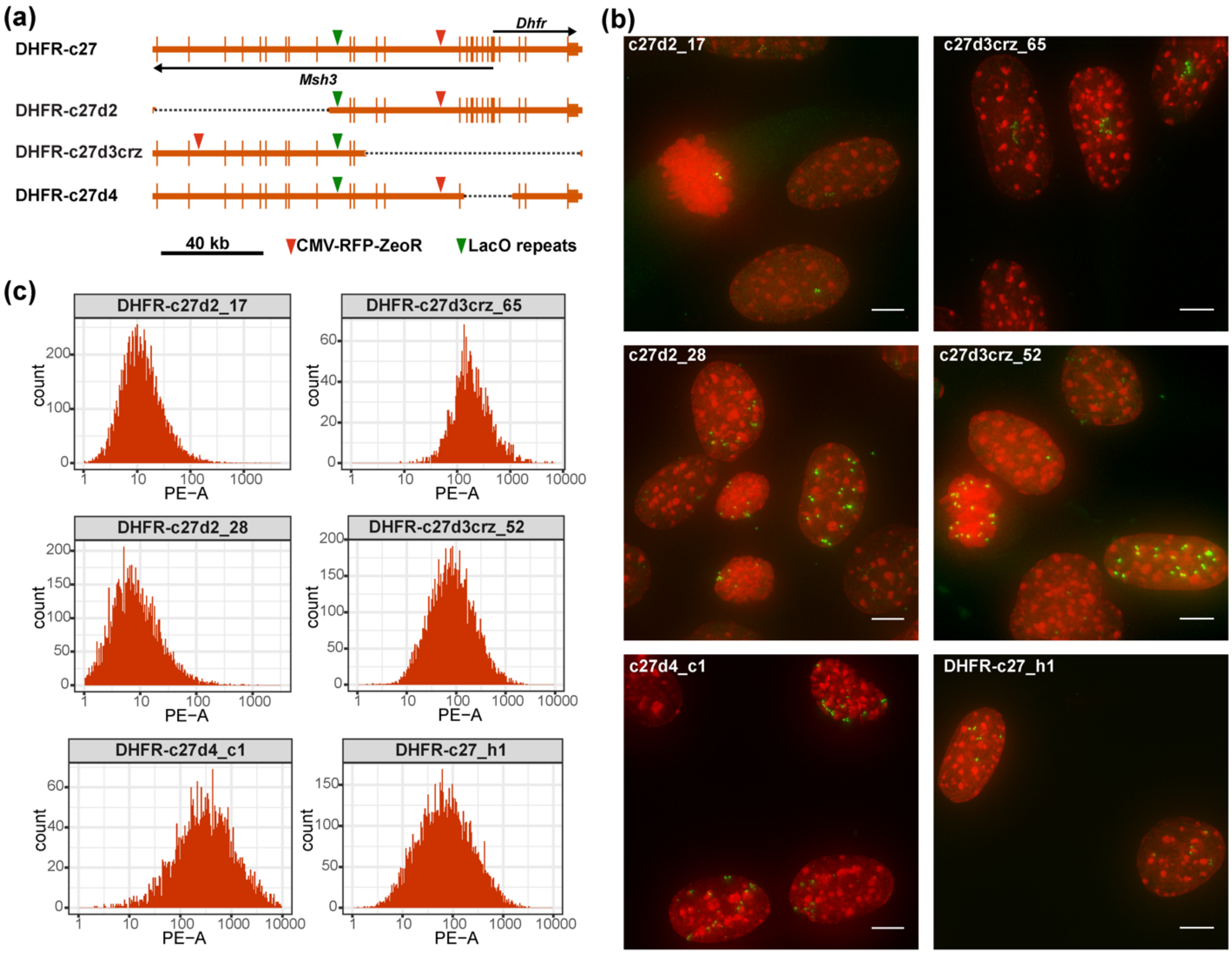
(a)
DHFR-c27
DHFR-c27d2
DHFR-c27d3crz
DHFR-c27d4
Dhfr
Msh3
40 kb
CMV-RFP-ZeoR
LacO repeats
(b)
c27d2_17
c27d3crz_65
c27d2_28
c27d3crz_52
c27d4_c1
DHFR-c27_h1
(c)
DHFR-c27d2_17
DHFR-c27d3crz_65
DHFR-c27d2_28
DHFR-c27d3crz_52
DHFR-c27d4_c1
DHFR-c27_h1
count
PE−A

Figure S9

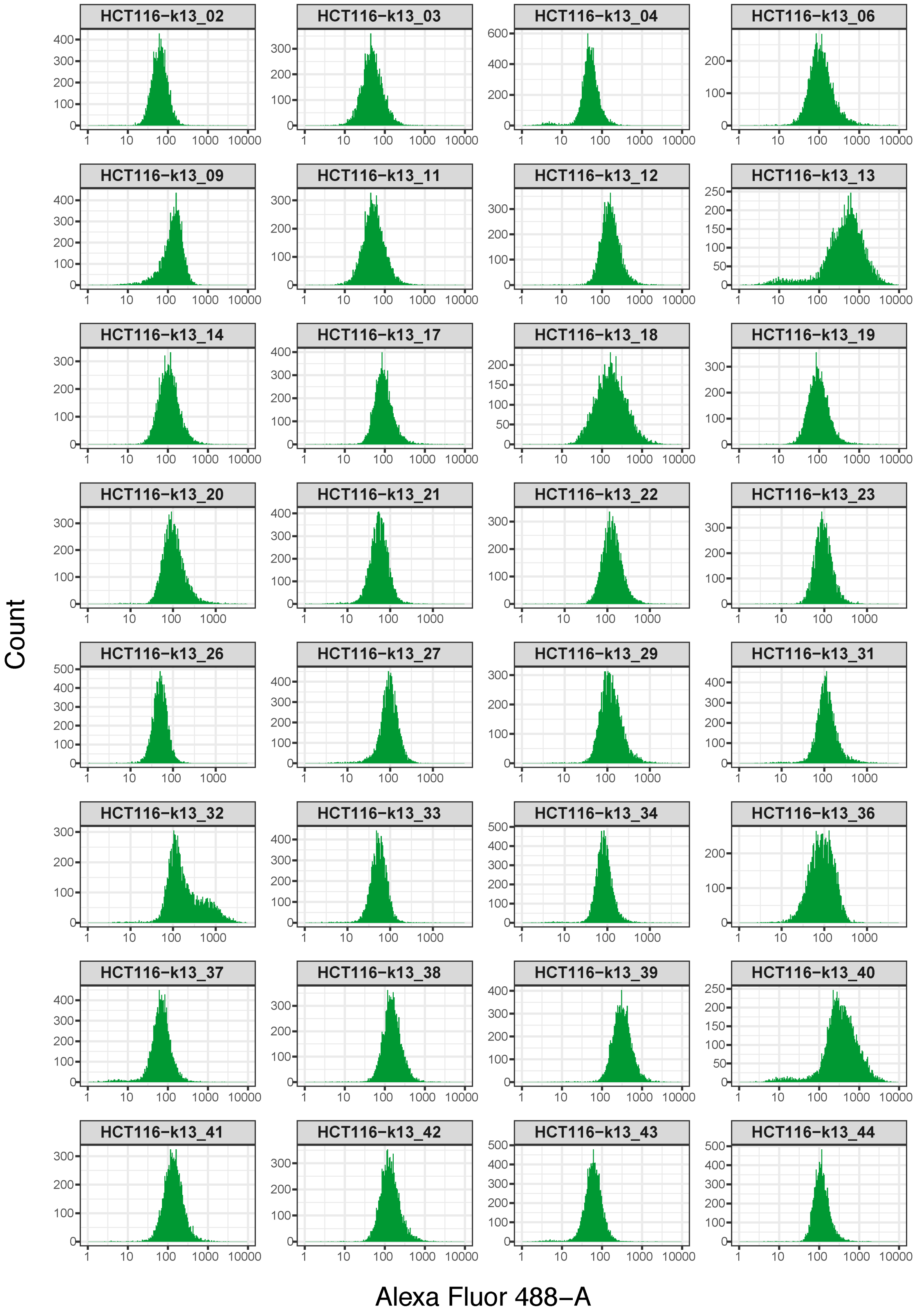
HCT116−k13_02
HCT116−k13_03
HCT116−k13_04
HCT116−k13_06
HCT116−k13_09
HCT116−k13_11
HCT116−k13_12
HCT116−k13_13
HCT116−k13_14
HCT116−k13_17
HCT116−k13_18
HCT116−k13_19
HCT116−k13_20
HCT116−k13_21
HCT116−k13_22
HCT116−k13_23
HCT116−k13_26
HCT116−k13_27
HCT116−k13_29
HCT116−k13_31
HCT116−k13_32
HCT116−k13_33
HCT116−k13_34
HCT116−k13_36
HCT116−k13_37
HCT116−k13_38
HCT116−k13_39
HCT116−k13_40
HCT116−k13_41
HCT116−k13_42
HCT116−k13_43
HCT116−k13_44
Count
Alexa Fluor 488−A

Figure S10

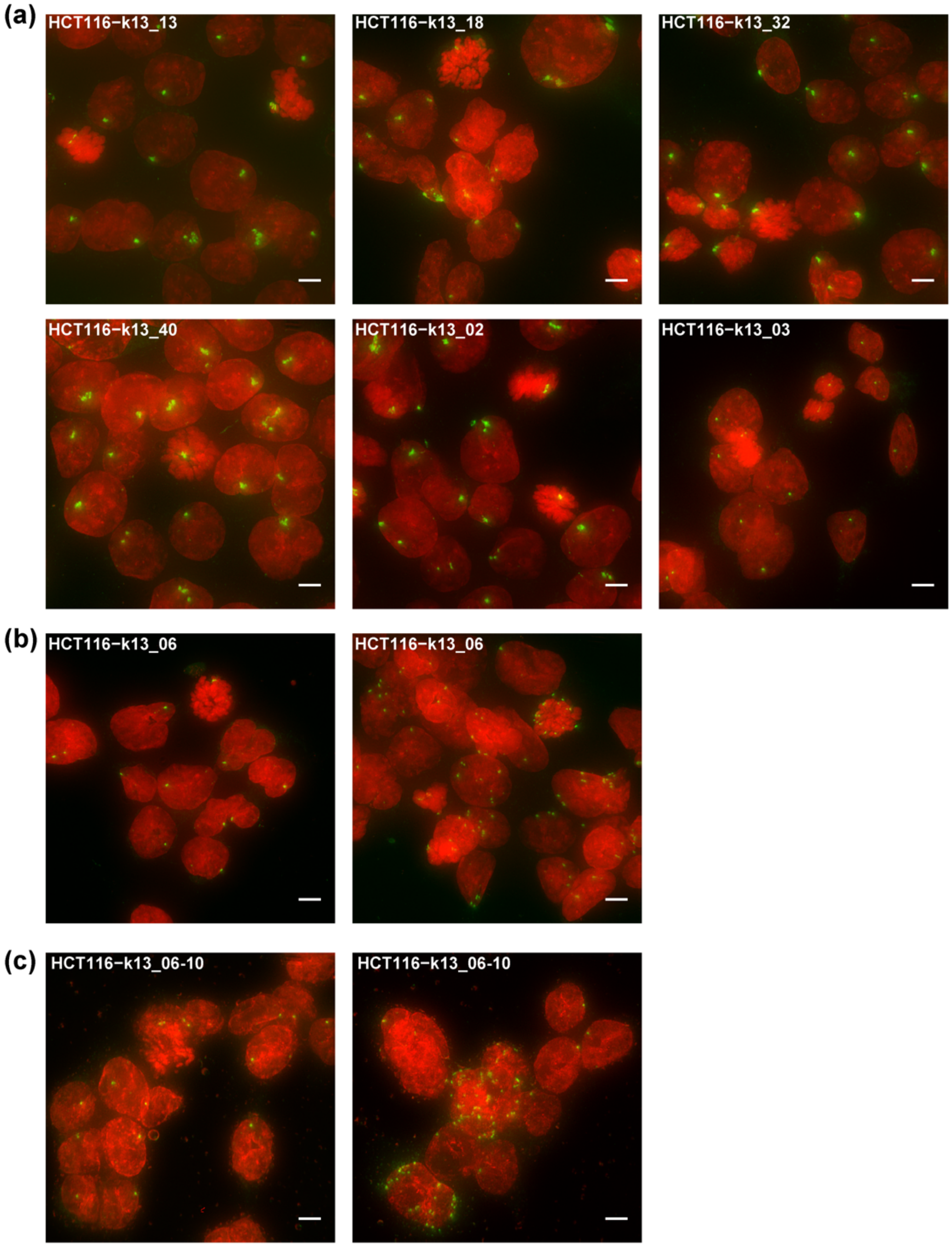
(a)
HCT116-k13_13
HCT116-k13_18
HCT116-k13_32
HCT116-k13_40
HCT116-k13_02
HCT116-k13_03
(b)
HCT116-k13_06
HCT116-k13_06
(c)
HCT116-k13_06-10
HCT116-k13_06-10

Figure S11

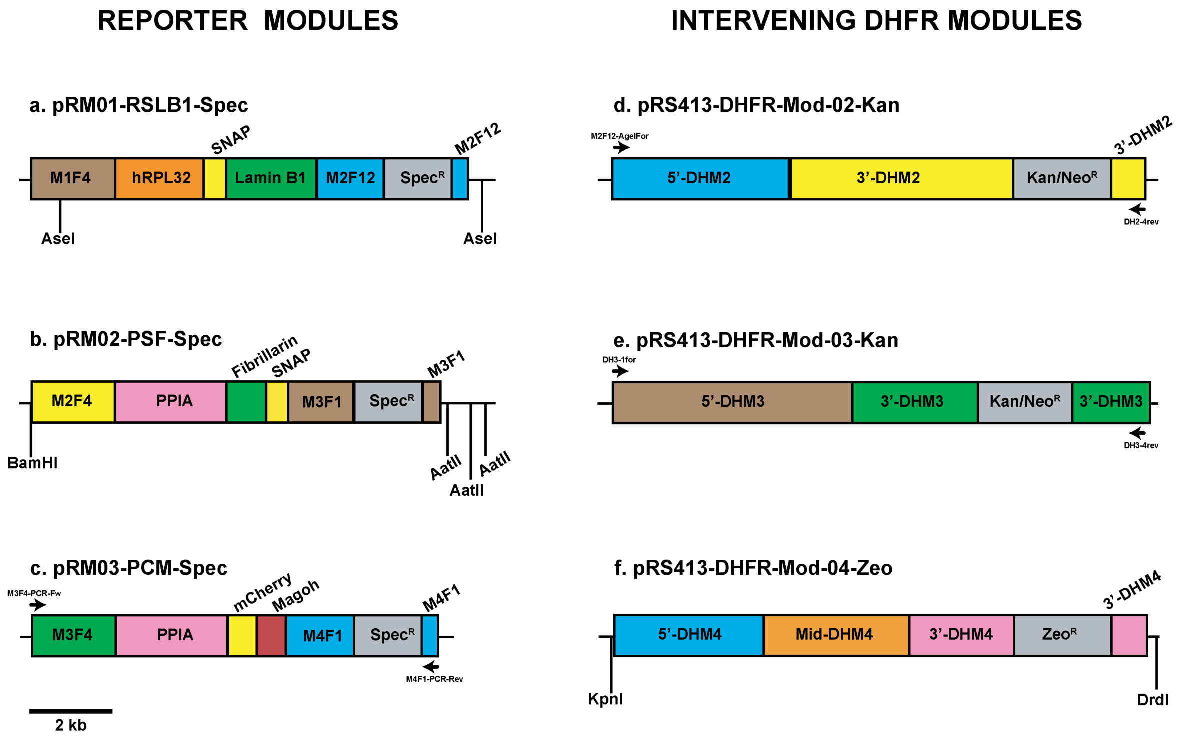
REPORTER MODULES
INTERVENING DHFR MODULES
a. pRM01-RSLB1-Spec
M1F4
hRPL32
SNAP
Lamin B1
M2F12
SpecR
M2F12
AseI
AseI
b. pRM02-PSF-Spec
M2F4
PPIA
Fibrillarin
SNAP
M3F1
SpecR
M3F1
BamHI
AatII
AatII
AatII
c. pRM03-PCM-Spec
M3F4-PCR-Fw
M3F4
PPIA
mCherry
Magoh
M4F1
SpecR
M4F1
M4F1-PCR-Rev
2 kb
d. pRS413-DHFR-Mod-02-Kan
M2F12-AgeIFor
5'-DHM2
3'-DHM2
Kan/NeoR
3'-DHM2
DH2-4rev
e. pRS413-DHFR-Mod-03-Kan
DH3-1for
5'-DHM3
3'-DHM3
Kan/NeoR
3'-DHM3
DH3-4rev
f. pRS413-DHFR-Mod-04-Zeo
5'-DHM4
Mid-DHM4
3'-DHM4
ZeoR
3'-DHM4
KpnI
DrdI

**Supplementary Figure S1.** GFP fluorescence histogram of representative "uniform" and "heterogeneous" expressing NIH 3T3 clones at day 0, 24, 60 and 96 without selection obtained by flow-cytometry. Gray- autofluorescence of untransfected cells; Green- GFP fluorescence of the indicated clones; x-axis- fluorescence; y-axis- cell number.

**Supplementary Figure S2.** DNA FISH over heterogeneously expressing clones transfected with DHFR-UG or HBB-UG BAC. (a-f) 3D DNA FISH over interphase nuclei from three DHFR-UG BAC (a-c) and three HBB-UG BAC (d-f) heterogeneous clones using BAC probes. Maximum-intensity projections are shown. (g-i) DNA FISH over mitotic spreads of one DHFR heterogeneous clone (g) and two HBB heterogeneous clones (h-i) using BAC probes. Red- DNA DAPI stain; Green- BAC FISH signal. Scale bar = 4 $\mu$m.

**Supplementary Figure S3.** Southern blot hybridization - using probes prepared from BAC DNA- of cellular DNA without enzyme digestion, or digested with increasing amount of S1 Nuclease, separated by PFGE. (a) Lane 1-2: NIH 3T3 cellular DNA; Lane 3-7: uniform clone HBB-UG-fD2; Lane 8-12: heterogenous clone HBB-UG-100d3; (b) *E. coli* carrying the HBB BAC.

**Supplementary Figure S4.** CNV analysis of the HBB-UG-100d3 clone. (a) Two FACS experiments for collecting cells with high (H1 and H2), and low (L) fluorescence subpopulations. x-axis- FITC channel intensity; y-axis- forward scatter; H1, H2, and Lsorting windows. (b) $\log_2$(ratio) of individual bins (dark gray dots) and the segment mean $\log_2$(ratio) (red lines) over the HBB BAC (pink highlight) in the H1 and H2 subpopulations of the HBB-UG-100d3 clone. (c) Scatter plot of segment mean $\log_2$(ratio) vs segment mean $\log_2$(normalized reads) of all segments of the H1 and H2 subpopulations of the HBB-UG-100d3 clone. Pink dots- segments belonging to the HBB BAC, including the HBB locus, UBC-GFP-ZeoR and the BAC vector; Black dots- genomic segments. (b-c) Blue dashed line: $\log_2$(estimated minimum copy number increase).

**Supplementary Figure S5.** Comparison of copy number fold increases of BAC regions (y-axis) in H1 and H2 relative to L subpopulations measured by CNV analysis versus by qPCR.

**Supplementary Figure S6.** $\log_2$(ratio) of individual bins (dark gray dots) and the segment mean $\log_2$(ratio) (red lines) over the UBC-GFP-ZeoR and the pBeloBAC11 BAC backbone, in the H1 and H2 subpopulations of the DHFR-UG-s3 and the HBB-UG-100d3 clones. Blue dashed line: $\log_2$(estimated minimum copy number increase).

**Supplementary Figure S7.** Promoter strength and BAC episome formation. (a) Summary of percentages of heterogeneously expressing clones and relative promoter strengths for different reporter constructs embedded in the DHFR BAC. (b) Average normalized RFP fluorescence of individual cell clones (y-axis) are plotted versus transgene copy number (x-axis) for EEF1α-mRFP-SV40-ZeoR (left) or RPL32-mRFP-

SV40-ZeoR (right) reporter constructs embedded in the DHFR BAC. Linear regression fits (black line) with y-intercepts set at 0 are shown with corresponding R-squared values and equations. Red circles- heterogeneous clones; Black circles- uniform clones. Bottom right: Number of clones analyzed and percentage of uniform clones. (c) Chromosomal locations of BAC transgene arrays within metaphase chromosomes of indicated clones as visualized by FISH using DHFR BAC probe (green) and DAPI staining (red). Scale bar = 5 μm.

**Supplementary Figure S8.** Episome formation of DHFR BACs with CMV-mRFP-SV40-ZeoR reporter gene insertions. (a) Schematics of the intact DHFR BAC (DHFR-c27) and three DHFR BAC deletions (DHFR-c27d2, -c27d3crz and -c27d4). Longer vertical bars- exons; shorter vertical bars- UTRs; arrows- direction of transcription; black dashed lines- deleted regions; green arrowhead- Lac operator repeats (LacO) insertion site; red arrowhead- CMV-mRFP-SV40-ZeoR insertion site. (b) Representative maximum-intensity projection images of clones with integrated BACs (c27d2_17, c27d3crz_65) and clones with episomal BACs (remaining clones). Red- DNA DAPI staining; Green- EGFP-LacI. Gamma = 0.5 was applied to the green channels of all images, and to the red channels of clone c27d2-17 and c27d3crz-52 images after projection. Scale bars = 5 μm. (c) mRFP fluorescence histogram of the clones in (b) obtained by flow-cytometry. x-axis- signal from PE channel; y-axis- cell number.

**Supplementary Figure S9.** GFP fluorescence histograms of HCT116 derived clones stably transfected with the 2207K13-UG BAC obtained by flow-cytometry. x-axis- signal from PE channel; y-axis- cell number.

**Supplementary Figure S10.** 3D DNA FISH over HCT116 derived clones stably transfected with the 2207K13-UG BAC using BAC probes. (a) Four clones with broad GFP-reporter fluorescence histograms (Supplementary Figure S9) show integrated BAC arrays (top 3 panels, left bottom panel), similar to clones with narrow GFP-reporter fluorescence histograms (middle and right bottom panels). (b-c) One clone with a narrow GFP-reporter fluorescence histogram showed a subpopulation of cells within the clonal population showing episomal BAC transgenes (b). Subcloning this clone identified a subclones which also contained a similar mixture of clones with integrated versus episomal forms of the the BAC transgenes (c). (a-c): Maximum intensity projections are shown. Gamma = 0.5 was applied to the green channels after projection. Red- DAPI staining; Green- FISH. Scale bars = 5 μm.

**Supplementary Figure S11.** Maps of Reporter and DHFR modules used for BAC-MAGIC. (a-c) reporter expression cassettes subcloned in the respective reporter recipient modules harboring SpecR selection marker (gray). (d-e) Schematics of the intervening DHFR modules harboring Kan/NeoR or ZeoR selection markers (gray). (a-e) Longer vertical bars represent the indicated restriction endonucleases used to generate recombineering fragments and arrows show the binding sites of primers used for amplification of recombineering fragments. See Methods for details of terminal regions corresponding to DHFR BAC homology regions. Scale bar = 2 kb.

**Supplementary Table S1.** List of primers used in this study.

**Supplementary Table S2.** Synthetic DNA fragment “RCS” sequence.
